# Glutamine transporters regulate prostate cancer radiosensitivity through NUPR1-mediated stress response

**DOI:** 10.1101/2025.04.22.650063

**Authors:** Uğur Kahya, Vasyl Lukiyanchuk, Ielizaveta Gorodetska, Matthias M. Weigel, Ayşe Sedef Köseer, Berke Alkan, Dragana Savic, Annett Linge, Steffen Löck, Mirko Peitzsch, Ira-Ida Skvortsova, Mechthild Krause, Anna Dubrovska

**Affiliations:** OncoRay - National Center for Radiation Research in Oncology, Faculty of Medicine and University Hospital Carl Gustav Carus, Technische Universität Dresden and Helmholtz-Zentrum Dresden-Rossendorf, Germany; Institute of Radiooncology - OncoRay, Helmholtz-Zentrum Dresden-Rossendorf (HZDR) Dresden, Germany; Department of Molecular Biology and Genetics, İzmir Institute of Technology, 35430 Izmir, Türkiye; Department of Therapeutic Radiology and Oncology, Medical University of Innsbruck, Innsbruck,Austria; EXTRO-Lab, Tyrolean Cancer Research Institute, Innsbruck, Austria; German Cancer Consortium (DKTK), partner site Dresden and German Cancer Research Center (DKFZ), Heidelberg, Germany; Department of Radiotherapy and Radiation Oncology, Faculty of Medicine and University Hospital Carl Gustav Carus, Technische Universität Dresden, Germany; National Center for Tumor Diseases (NCT), partner site Dresden: German Cancer Research Center (DKFZ), Heidelberg; Faculty of Medicine and University Hospital Carl Gustav Carus, Technische Universität Dresden, and Helmholtz-Zentrum Dresden-Rossendorf (HZDR), Dresden, Germany; Institute for Clinical Chemistry and Laboratory Medicine, University Hospital and Faculty of Medicine Carl Gustav Carus, Technische Universität Dresden, Dresden, Germany

**Keywords:** prostate cancer, radiation therapy, Glutamine transporters, GLS, NUPR1, oxidative stress

## Abstract

**Background:** Metabolic and stress response adaptations in prostate cancer (PCa) mediate tumor resistance to radiation therapy (RT). Our study investigated the roles of glutamine (Gln) transporters *SLC1A5, SLC7A5*, and *SLC38A1* in regulating *NUPR1*-mediated stress response, PCa cell survival, metabolic reprogramming, and response to RT.

**Methods:** The radiosensitizing potential of GLS inhibition with CB-839 was analyzed in prostate cancer xenograft models. The level of gene expression was analyzed by RNA sequencing and RT-qPCR in the established cell lines or patient-derived tumor and adjacent non-cancerous tissues. Phosphoproteomic analysis was employed to identify the underlying signaling pathways. The publicly available PCa patient genesets, and a geneset for the patients treated with RT were analyzed by SUMO software. The key parameters of mitochondrial functions were measured by Seahorse analysis. Analysis of the general oxidative stress level and mitochondrial superoxide detection were conducted using flow cytometry. γH2A.X foci analysis was used to assess the DNA double strand break. Relative cell sensitivity to RT was evaluated by radiobiological clonogenic assays. Aldefluor assay and sphere-forming analysis were used to determine cancer stem cell (CSC) phenotype.

**Results:** Depletion of these transporters led to reduced cell viability, altered ROS levels, and enhanced radiosensitivity in PCa cell lines. Functional assays revealed that targeting these transporters decreases CSC properties, impairs cell cycle progression, and deregulates mitochondrial energy metabolism. Our findings indicate that the Gln transporters mediate the adaptation of tumor cells to nutrient stress, while *NUPR1* promotes cellular survival under metabolic and genotoxic stress. Targeting *SLC1A5*, *SLC7A5*, *SLC38A1*, and *NUPR1* radiosensitizes PCa by disrupting metabolic adaptations and stress responses. Besides, CB-839, a glutaminase (GLS) inhibitor, combined with RT, demonstrated a synergistic effect with radiotherapy *in vivo*, significantly delaying tumor growth. We found that *GLS* gene expression levels are significantly associated with clinical outcomes in PCa patients treated with RT.

**Conclusions:** Our work underscores the role of Gln transporters and the NUPR1-mediated stress response induced by Gln deficiency in PCa cell survival, stemness, mitochondrial functions and radioresistance. Our findings provide a potential therapeutic *in vivo* strategy to enhance the efficacy of RT and improve treatment outcomes for PCa patients.

## Background

Prostate cancer (PCa) is one of the most common cancers in men globally (1). Radiotherapy (RT) is an essential treatment option for PCa, together with surgery and chemotherapy (2, 3). Despite advances in early detection and localized treatment, resistance to conventional therapies such as RT remains a significant clinical challenge (4). Cancer is a metabolic disease, and tumor development is driven by metabolic reprogramming (5, 6). This hallmark of cancer enables tumor cells to adapt to therapeutic stress and sustain proliferation. Thereby, tumorigenesis is linked to the accumulation of genetic mutations and epigenetic modifications that promote uncontrolled proliferation, consequently elevating the demand for nutrients in tumors (7). As a result, tumor cells enhance the uptake of glucose and amino acids to meet the increased energetic and biosynthetic requirements associated with rapid proliferation and adaptation to environmental stresses (8, 9).

Glutamine (Gln) is the most abundant amino acid in plasma, a critical nutrient for cellular growth, energy production, biosynthesis of proteins and nucleotides, and maintenance of redox balance (10). Cancer cells often exhibit increased Gln uptake and metabolism, a phenomenon known as “Gln addiction” (11–13). Gln is vital for cancer cells, as it fuels the tricarboxylic acid (TCA) cycle to generate ATP through mitochondrial oxidative phosphorylation (OXPHOS). Glutaminolysis not only provides the intermediates for the TCA cycle but also fuels lactate production and helps to recycle nicotinamide adenine dinucleotide (NADH) and flavin adenine dinucleotide (FADH2), which are essential reducing equivalents used by the electron transport chain (ETC) (13–18). Gln contributes to the oxidative stress defense by mediating the production of a key scavenger of the reactive oxygen species, glutathione (GSH), from Gln-derived glutamate (Glu). GLS is one of the enzymes catabolizing Gln to Glu. Elevated GLS activity has been associated with tumor growth, poor prognosis, and therapy resistance (19–21). Amino acid availability supports redox balance by regulating the mechanistic target of rapamycin (mTOR)-driven antioxidant defense, endoplasmic reticulum (ER) stress, and unfolded protein response (UPR) pathways (22–24). Gln uptake and catabolism regulate the intracellular pool of α-Ketoglutarate (α-KG), and α-KG -driven epigenetic reprogramming (18). Recent studies also highlight epigenetic mechanisms that help cells counter oxidative stress induced by amino acid uptake and transporter expression (7, 25, 26).

Amino acid transporters are essential for the uptake of Gln and other amino acids to regulate intracellular amino acid homeostasis, redox balance, and cellular growth (7). Among these, *SLC1A5* (also known as ASCT2), *SLC7A5* (LAT1) and its heavy chain *SLC3A2* (CD98hc), and *SLC38A1* (SNAT1) are highly expressed in various cancers and have been implicated in tumor progression, therapeutic resistance and clinical outcome (7, 27–39). *SLC1A5* primarily mediates Gln uptake, while *SLC7A5* functions as a bidirectional transporter facilitating the exchange of intracellular Gln for extracellular essential amino acids, including leucine (40–43). This exchange activates the mTOR signaling pathway, promoting protein synthesis and cell growth (44). *SLC38A1* contributes to the uptake of not only Gln but also other neutral amino acids, supporting nucleotide and protein biosynthesis (7, 45). Moreover, transcription factor *MYC* regulates the expression of genes involved in Gln metabolism, including GLS and amino acid transporters (12, 46). MYC-driven upregulation of these metabolic genes enhances Gln uptake and utilization, supporting the anabolic demands of rapidly proliferating cancer cells (12, 46–48). Moreover, metabolic stress induced by Gln deprivation or GLS inhibition can activate stress response signaling, such as the UPR pathway and the integrated stress response (ISR) (44, 49, 50).

Nuclear protein 1 (*NUPR1*), a stress-inducible transcriptional regulator, is upregulated in various tumor entities and is associated with therapy resistance, promoting cell survival under metabolic and genotoxic stress conditions (51–55). We and others have demonstrated that targeting Gln metabolism can sensitize cancer cells to RT and other therapies (19, 21, 56–59). However, the role of amino acid transporters in modulating PCa cell response to RT and the underlying mechanisms involving stress response proteins like *NUPR1* remain insufficiently understood. Elucidating these mechanisms could identify novel therapeutic targets to overcome resistance and improve treatment outcomes.

In this study, we employed metabolic analyses, gene expression profiling, and functional assays to investigate how the amino acid transporters *SLC1A5*, *SLC7A5*, and *SLC38A1*, along with the stress response protein *NUPR1*, regulate PCa cell survival, metabolism, stemness, and response to RT. Our findings demonstrate that targeting these Gln transporters and *NUPR1* disrupts Gln uptake and metabolism, consequently enhancing PCa radiosensitivity. The inhibition of a glutaminase (GLS) with CB-839, the only clinically approved inhibitor of glutaminolysis, demonstrated a synergistic effect with radiotherapy *in vivo*. Using patient-derived tumor and adjacent non-cancerous tissues, we found that expression of the *SLC1A5*, *SLC7A5*, and *SLC38A1*, and *NUPR1* is a rescue mechanism induced by GLS inhibition. This work elucidates the factors driving therapeutic resistance, providing a rationale for novel combination therapies in PCa.

## Methods

### Cell lines

PCa cell lines DU145, PC3, and LNCaP were obtained from the American Type Culture Collection (ATCC). As previously described (21, 60), the radioresistant (RR) cell lines were generated by *in vitro* selection using fractionated X-ray irradiation. Cells were cultured in Dulbecco’s Modified Eagle’s Medium (DMEM; Sigma-Aldrich) or RPMI-1640 medium (Sigma-Aldrich), supplemented with 10% fetal bovine serum (FBS; Capricorn Scientific), 2 mM L-Gln (Sigma-Aldrich), 10 mM HEPES buffer (Sigma-Aldrich), 1 mM sodium pyruvate (Sigma-Aldrich), and 1× MEM non-essential amino acids (MEM-NEAA) (Sigma-Aldrich). Cells were maintained at 37°C in a humidified atmosphere with 5% CO_2_ and passaged at 70-90% confluence.

For Gln starvation experiments, cells were cultured in DMEM or RPMI-1640 medium supplemented with 10% dialyzed FBS (Sigma-Aldrich) and 10 mM HEPES buffer (Sigma-Aldrich). Control cells were cultured in DMEM or RPMI-1640 medium supplemented with 10% dialyzed FBS, 2 mM L-Gln (Sigma-Aldrich), and 10 mM HEPES buffer (Sigma-Aldrich). All cell lines were tested negative for mycoplasma and genotyped using microsatellite polymorphism analysis.

### Clinical specimens

The formalin-fixed paraffin-embedded (FFPE) tumor tissues of patients (n=67) with intermediate- or high-risk localized PCa who were treated with curatively-intended, definitive RT at the Department of Radiotherapy and Radiation Oncology, University Hospital Carl Gustav Carus and Faculty of Medicine, Dresden, were analyzed by a whole transcriptomics using the HTA 2.0 Array (Affymetrix). Patient clinical characteristics were described previously (21). A freedom from PSA relapse was used as clinical endpoint. Survival curves were analyzed by the Kaplan–Meier method.

### siRNA-mediated gene silencing

A total number of 200,000 DU145 and PC3 cells, as well as 300,000 LNCaP cells, were seeded into 6-well plates. After 24 hours of incubation, DU145 and PC3 cells were transfected with pooled siRNA constructs targeting the gene of interest. For transfection, 9 µL of Invitrogen™ Lipofectamine™ RNAiMAX was mixed with 150 µL of Opti-MEM. In a separate tube, 40 pmol of siRNA was mixed with 150 µL of Opti-MEM. The siRNA-Opti-MEM mixture was combined with the Lipofectamine-Opti-MEM mixture and gently added dropwise to the wells after 5-minute incubation. Following the addition of transfection reagents, cells were incubated at 37°C in a CO_2_ incubator for 24 hours before being used in subsequent experiments. LNCaP cells were transfected 48 hours after seeding under the same conditions. Cells transfected with pooled non-targeting siRNA (scrambled siRNA or siSCR) were the negative control in all knockdown experiments. 24 hours after siRNA-mediated gene silencing, cells were harvested using 0.25% trypsin-EDTA solution (Gibco) and centrifuged at 1,000 rpm for 5 minutes. The supernatant was discarded, and the cells were resuspended in a complete DMEM or RPMI-1640 medium. The cells were then counted using a Neubauer chamber and seeded accordingly. The siRNA duplexes were synthesized by Eurogentec. Gene-silencing was validated by RT-qPCR. The siRNA sequences are listed in Supplementary Table 1.

### shRNA-mediated gene silencing

A total number of 200,000 DU145 and PC3 cells, as well as 300,000 LNCaP cells, were seeded into 6-well plates. After 24 hours of incubation, cells were transfected with pLKO.1 puro vector constructs expressing shRNA against human *SLC1A5*, *SLC7A5*, and *SLC38A1* or nonspecific control shRNA (shControl) using Lipofectamine 2000 transfection reagent (Thermo Scientific) according to the manufacturer’s instructions. 48 h after transfection, cells were selected with puromycin at 1 µg/ml concentration for a week and further selection with puromycin at 5 µg/ml concentration applied for another week. The list of shRNA expression vectors is provided in Supplementary Table 2.

### Clonogenic cell survival assay

Cells were seeded in triplicates into 6-well plates at a density of 2,000 cells per well. The following day, the plates were irradiated with 0 Gy, 2 Gy, 4 Gy, or 6 Gy of X-rays (Yxlon Y.TU 320; 200 kV X-rays, dose delivery rate 1.3 Gy/min at 20 mA, filtered with 0.5 mm Cu). After irradiation, the cells were incubated at 37°C in a CO_2_ incubator for 7 to 10 days to allow colony formation.

Following the incubation period, the plates were fixed with 10% formaldehyde in PBS for 30 minutes and stained with 0.05% crystal violet solution for 30 minutes at room temperature. The plates were then air-dried, and colonies containing more than 50 cells were manually counted using a stereomicroscope (Zeiss). The plating efficacy (PE) at 0 Gy and survival fraction (SF) were calculated as described earlier (120). Results were presented as survival fractions depicted on a logarithmic scale and plotted against applied X-ray doses.

### Cell viability assay

Cells were seeded into 96-well plates at a density of 10,000 for DU145 and 20,000 for LNCaP cells per well, respectively. The following day, the medium was carefully aspirated using a multichannel aspirator, and cells were resuspended in 80 µL of a 1:1 mixture of PBS and CellTiter-Glo® reagent (Promega). The plates were shaken at room temperature for 30 minutes to ensure complete cell lysis. After confirming cell lysis, the lysate was transferred to an opaque-walled white multiwell plate, and luminescence was measured using the SpectraMax® iD3 microplate reader (Molecular Devices).

For IC50 calculations of ZZW-115, cells were seeded into 96-well plates at a density of 10,000 for DU145 and 20,000 for LNCaP cells per well, respectively. The following day for DU145 cells and 48 hours after plating for LNCaP cells, serial dilution of ZZW-115 from 30µM to 0µM were applied to wells sequentially. 48 hours after the drug treatment, the medium was carefully aspirated using a multichannel aspirator, and cells were resuspended in 80 µL of a 1:1 mixture of PBS and CellTiter-Glo® reagent (Promega). The plates were shaken at room temperature for 30 minutes to ensure complete cell lysis. After confirming cell lysis, the lysate was transferred to an opaque-walled white multiwell plate, and luminescence was measured using the SpectraMax® iD3 microplate reader (Molecular Devices).

### Cell Cycle Distribution Analysis

Cell cycle distribution was analyzed in DU145 and LNCaP cells following siRNA-mediated gene silencing and subsequent irradiation with 6 Gy X-rays (Yxlon Y.TU 320; 200 kV X-rays, dose delivery rate 1.3 Gy/min at 20 mA, filtered with 0.5 mm Cu). Cells were seeded in 6-well plates and harvested post-irradiation. Harvested cells were washed with Flow Cytometry buffer (1X DPBS, 1 mM EDTA, 25 mM HEPES, 3% FBS) and incubated with 10 µg/mL Hoechst 33342 DNA dye (Invitrogen) diluted in Flow Cytometry buffer. The cells were incubated for 45 minutes at 37°C.

After incubation, the reaction was halted by placing the samples on ice. Immediately prior to flow cytometric analysis, 2 µg/mL of the live-dead marker 7-Aminoactinomycin (7-AAD) (Sigma-Aldrich) was added to the samples. Flow Cytometry was performed using a BD FACSCelesta Flow Cytometer (BD), and data analysis was carried out with FlowJo software. Sham-irradiated cells were used as controls for the analysis.

### Cell Death Assay

Following the manufacturer’s instructions, cell death was assessed using the eBioscience™ Annexin V Apoptosis Detection Kit (Thermo Fisher Scientific) after siRNA-mediated gene silencing and irradiation with 6 Gy X-rays (Yxlon Y.TU 320; 200 kV X-rays, dose delivery rate 1.3 Gy/min at 20 mA, filtered with 0.5 mm Cu). Briefly, all cells were harvested using Accutase, washed once with PBS, and then washed again with 1X Annexin V buffer. 100,000 cells were resuspended in 100 μL of 1X Annexin V buffer and stained with FITC-conjugated Annexin V for 15 minutes at room temperature. After incubation, cells were washed once with 1X Annexin V buffer and resuspended in 200 μL of the same buffer.

Subsequently, 5 μL of propidium iodide (PI, Thermo Fisher Scientific) was added to the cell suspension. The fluorescence of the cells was then measured using a BD FACSCelesta flow cytometer, and data were analyzed using FlowJo software. Sham-irradiated cells were used as controls.

### Sphere formation assay

Harvested cells were washed once with PBS and resuspended as single cells in Mammary Epithelial Cell Growth Medium (MEBM; Lonza) supplemented with B27 (Invitrogen), 4 μg/mL insulin (Sigma-Aldrich), 1 mM L-Gln (Sigma-Aldrich), 20 ng/mL epidermal growth factor (EGF; Peprotech), and 20 ng/mL basic fibroblast growth factor (FGF; Peprotech). Cells were plated into 24-well ultra-low attachment plates (Corning) at a density of 5,000 cells per well for DU145 and PC3 and 2,000 cells per well for LNCaP in 1 mL of MEBM medium. One week later, 1 mL of MEBM medium was added to each well, and cell clumps were gently disaggregated by pipetting. Two weeks after the initial cell plating, the plates were scanned using the Celigo S Imaging Cell Cytometer (Brooks). During image analysis, spheres were distinguished from cell aggregates by a diameter ≥ 50 µm and a roundish shape. Sphere size was determined from the images using Fiji/ImageJ software (61).

### Fluorescence microscopy of γH2A.X foci slides

For γH2A.X foci staining, cells were plated onto 8-well Millicell® EZ chamber slides (Merck Millipore) at a density of 25,000 and 30,000 cells per well for DU145 and LNCaP, respectively, and incubated at 37°C in a CO₂ incubator. The following day, cells were irradiated with 4 Gy X-rays (Yxlon Y.TU 320; 200 kV X-rays, dose delivery rate 1.3 Gy/min at 20 mA, filtered with 0.5 mm Cu). Sham-irradiated cells were used as controls. Twenty-four hours after irradiation, cells were fixed with 3.7% formaldehyde (Thermo Fisher Scientific) for 30 minutes at 37°C. After washing the samples three times with PBS, the samples were permeabilized with 0.25% Triton X-100 (Sigma-Aldrich) for 7 minutes at room temperature and washed again with PBS. The samples were then blocked with 10% BSA (Fisher Scientific) in PBS at 37°C for one hour.

Subsequently, the cells were incubated overnight at 4°C with the primary anti-γH2A.X mouse antibody (Merck Millipore), diluted as 1:400 in 3% BSA/PBS. The following day, the samples were washed 10 times with PBS and incubated with the secondary AlexaFluor™ 488 goat anti-mouse antibody, diluted 1:500 in 3% BSA/PBS, for one hour at room temperature in the dark. After additional PBS washes, nuclear staining was performed with 1 µg/mL DAPI for 5 minutes. After removing excess DAPI by washing the slides with PBS, Mowiol solution was added to the slides covered with glass coverslips. The images were taken with a confocal Leica SP5 microscope or an LSM 980 with Airyscan 2 (Zeiss). Images were analyzed using Fiji/ImageJ software to count the number of foci per nucleus (61).

### ALDEFLUOR assay

Aldehyde dehydrogenase (ALDH) activity was analyzed using the Aldefluor™ assay (Stem Cell Technologies) following the manufacturer’s protocol. Briefly, cells were detached using trypsin (Sigma-Aldrich), washed with PBS, and resuspended in Aldefluor buffer. To identify ALDH-positive cells, the cells were incubated with diethylaminobenzaldehyde (DEAB), a specific ALDH inhibitor, at a 1:50 dilution, serving as a negative control. Both control and test samples were stained with the Aldefluor reagent at a 1:200 dilution and incubated at 37°C for 30 minutes. Gating was performed based on the DEAB control to determine the ALDH-positive cell population.

### RNA isolation, cDNA synthesis, and RT-qPCR

RNA from PCa cells was isolated using the NucleoSpin RNA Plus kit (Macherey-Nagel) following the manufacturer’s instructions. Reverse transcription was performed using the PrimeScript™ RT reagent Kit (Takara) according to the manufacturer’s protocol. The RNA volume used for reverse transcription was adjusted across all samples to achieve a unified RNA concentration.

Quantitative real-time polymerase chain reaction (qRT-PCR) was carried out using the TB Green Premix Ex Taq II (Takara Bio Inc) according to the manufacturer’s protocol, with a total reaction volume of 20 μL. The qPCR cycling program was conducted on a StepOnePlus system (Applied Biosystems) with the following settings: 94°C for 2 minutes, followed by 40 cycles of 94°C for 15 seconds, 58°C for 60 seconds, and 72°C for 60 seconds, concluding with a melt curve analysis performed by increasing the temperature to 95°C in steps of 0.3°C.

All experiments were performed with at least three technical replicates. The expression levels of ACTB and RPLP0 mRNA were used as housekeeper genes for data normalization. The primers used in the study were purchased from Eurofins Genomics Germany GmbH and are listed in Supplementary Table 3.

### Western Blotting

Cells were harvested by scraping into RIPA lysis buffer on ice. Lysates were centrifuged at 13000 rpm for 10 min, and the supernatant was transferred to the new tubes. Measurement of the total protein concentration was done by using BCA assay kit (Pierce) in 96-well plate format. Samples were mixed with 4x Laemmli Sample Buffer (Bio-Rad) containing dithiothreitol (DTT) at a final concentration of 10 mM. Samples were denaturated at 96°C for 5 min, cell lysates were loaded onto NuPAGE Bis-Tris 4-12% acrylamide gels (Thermo Fischer Scientific) along with prestained protein ladder PageRuler (10-180 kDa) (Thermo Fisher Scientific) and run at 120 V. Proteins were transferred to the Protran® nitrocellulose membrane (Whatman) for 2 hours at 250 mA with cooling. The transfer quality was confirmed with Ponceau S solution (Sigma). Membranes were blocked in 5% BSA in PBS for 1 h at room temperature and incubated with primary antibodies overnight on the rocker shaker at 4°C. Primary antibodies were used at concentrations recommended by the manufacturer. Next day membranes were washed 5 times for 10 min in 0.05% PBS/Triton X-100 solution, and corresponding secondary antibodies conjugated with horseradish peroxidase (GE Healthcare) were added. Membranes were incubated with secondary antibodies 1 hour on the shaker at room temperature, then washed 3 times with PBS/Triton X-100 solution, covered with freshly prepared SuperSignal West Dura Extended Duration Substrate (Thermo Fischer Scientific), and exposed to the LucentBlue X-ray film (Advansta Inc.). Antibodies used for Western blot analysis are listed in Supplementary Table 4.

### Seahorse analysis

For seahorse analysis, 24h after transfection, cells were seeded in triplicates into the Seahorse XFp Cell Culture Miniplates (Agilent Technologies) in a growth medium and incubated overnight at 37°C and 5% CO_2_. The Seahorse XFp sensor cartridges were hydrated at 37°C with Seahorse XF calibrant in a non-CO_2_ incubator overnight. On the day of the experiment, the cell medium was changed to the corresponding assay medium: Seahorse XF RPMI Medium, pH 7.4 (LNCaP) or Seahorse XF DMEM medium, pH 7.4 (DU145), supplemented with 10 mM of glucose, 2 mM of l-Gln, and 1 mM of sodium pyruvate. Cells were incubated for 1 h prior to the assay at 37°C in a non-CO_2_ incubator.

The Seahorse XF Mito Stress Test was performed according to the manufacturer’s protocol. After 18 min of basal condition measurements, serial injections of 1.5 µM oligomycin, 1 µM FCCP, and 0.5 µM rotenone/antimycin A (final well concentrations) followed. Oxygen consumption rate (OCR) were measured using the Agilent Seahorse XFp Analyzer (Agilent Technologies, California, USA). Background recordings were acquired from wells with medium only (without cells) and subtracted. The obtained results were normalized to the final number of cells at each well. Following the assay, cells were stained with Hoechst solution, images of whole wells were taken by a Lionheart FX automated microscope (Agilent BioTek), and cell numbers were assessed using the open-access ImageJ/Fiji software (61). Data analysis and reporting of mitochondrial function were conducted using Seahorse Wave Desktop software and the Seahorse XF Cell Mito Stress Test report generator.

### RNA-Sequencing and gene enrichment analysis

The RNA-Seq dataset was processed using the Statistical Utility for Microarray and Omics (SUMO) software (https://angiogenesis.dkfz.de/oncoexpress/software/sumo/). First, genes expressed at noise levels were filtered out to reduce background interference. The remaining data were then quantile normalized across all samples to ensure comparability. Gene expression values were further normalized using the median and transformed to the log2 scale to stabilize variance and normalize the distribution. Significantly deregulated genes predicted as off-targets by the DSIR web tool (http://biodev.cea.fr/DSIR/DSIR.html) were excluded from the dataset. The remaining deregulated genes were analyzed to assess their enrichment or depletion between different treatment groups, comparing observed differences to those expected by random chance. log2 fold change (log2FC) of genes in the *SLC1A5* siRNA, *SLC7A5* siRNA, and *SLC38A1* siRNA conditions was calculated by subtracting the log2 values of the Scr siRNA condition from those of each experimental condition. Then, gene sets derived from “msigdbr” package of R Programming were used in Gene Set Enrichment Analysis (GSEA); “fgsea” package of R programming was used to perform GSEA, and the results were visualized using the “ggplot2” package.

### Analysis of the patient cohort data

The Publicly available datasets, including TCGA PRAD and MSKCC, were downloaded from cBioPortal for Cancer Genomics (https://www.cbioportal.org/). The data were analyzed using SUMO software. For Kaplan-Meier survival analysis, biochemical recurrence-free survival time was determined using the “Days to PSA” and “Days to biochemical recurrence first” data provided in the datasets. Patient groups were stratified based on the optimal cut-off point identified through a scan procedure.

### Ingenuity Pathway Analysis (IPA)

Differentially expressed genes were identified by using SUMO software and were analyzed using Ingenuity Pathway Analysis (IPA, Qiagen-https://digitalinsights.qiagen.com/IPA). Gene expression data were uploaded to the IPA platform, including log2 fold changes and associated p-values. The data were mapped to corresponding gene objects in the Ingenuity Knowledge Base.

Canonical pathway analysis, upstream regulator analysis, and network analysis were performed. Pathways, molecular functions, and biological processes that were significantly enriched among the differentially expressed genes were identified. The significance of canonical pathways was assessed by Fisher’s exact test, and z-scores were used to predict the activation or inhibition states of pathways or upstream regulators. The analysis focused on identifying key regulatory pathways, molecular interactions, and biological networks affected by the different treatments. Only pathways with a p-value < 0.05 and relevant Log2FC cutoff (≥ |0.2|) were considered significantly activated or inhibited.

### Mass spectrometry-based analysis of Krebs cycle metabolites

TCA cycle metabolites in conjunction with amino acids Gln, glutamic acid, asparagine and aspartic acid as well as lactate, 2-hydroxyglutarate and arginine-succinate were analyzed by liquid chromatography-tandem mass spectrometry (LC-MS/MS) as described elsewhere (62).

### Analysis of the oxidative stress

General oxidative stress levels in DU145 and LNCaP cells were assessed following siRNA-mediated gene silencing and exposure to 6 Gy X-ray irradiation (Yxlon Y.TU 320; 200 kV X-rays, dose rate 1.3 Gy/min at 20 mA, filtered with 0.5 mm Cu). Cells were seeded in 6-well plates, and 24 hours post-irradiation, they were harvested using Accutase (StemCell Technologies). The harvested cells were washed with Flow Cytometry buffer (1X DPBS, 1 mM EDTA, 25 mM HEPES, 3% FBS). Approximately 200,000 cells in 200 µL of flow cytometry buffer were stained with 1 µM of the oxidative stress indicator 2’,7’-dichlorodihydrofluorescein diacetate (CM-H2DCFDA, Invitrogen) for 30 minutes at 37°C.

After staining, cells were washed once and re-suspended in a flow cytometry buffer. 2 µg/mL of 7-amino actinomycin D (7-AAD, Sigma-Aldrich) was added to the cell suspension to exclude dead cells. The green fluorescence intensity, indicative of reactive oxygen species (ROS) production, was measured using a BD FACSCelesta Flow Cytometer (BD). Data analysis was conducted with FlowJo software. Sham-irradiated cells served as controls in the analysis.

### Mitochondrial Superoxide Detection

Mitochondrial superoxide production in DU145 and LNCaP cells was evaluated following siRNA-mediated gene silencing and exposure to 6 Gy X-ray irradiation (Yxlon Y.TU 320; 200 kV X-rays, dose rate 1.3 Gy/min at 20 mA, filtered with 0.5 mm Cu). Cells were seeded in 6-well plates and harvested 24 hours post-irradiation using Accutase (StemCell Technologies). After harvesting, cells were washed with Flow Cytometry buffer (1X DPBS, 1 mM EDTA, 25 mM HEPES, 3% FBS).

Approximately 200,000 cells were resuspended in 200 µL of Flow Cytometry buffer and stained with 0.5 µM MitoSOX™ Mitochondrial Superoxide Indicator Red (Thermo Fisher Scientific) for 30 minutes at 37°C, protected from light. Following staining, cells were washed once with Flow Cytometry buffer and resuspended. 1 µg/mL of DAPI (Thermo Fisher Scientific) was added to the cell suspension to exclude dead cells.

### Tumor growth in mice xenograft models

The animal experiments were conducted in the OncoRay facility in Dresden, following institutional guidelines and German animal welfare regulations (protocol number TVV21/2022). Five-week-old male Rj: NMRI-Foxn1^nu/nu^ mice were ordered from Janvier-Labs (France), and experiments were conducted using 8- to 10-week-old male NMRI (nu/nu) mice. Whole-body irradiation was administered to the animals one day before tumor transplantation at 4 Gy (200 kV X-rays, 0.5 mm Cu filter, 1.3 Gy/min) for immunosuppression. Subcutaneous (s.c.) xenograft tumors were generated by injecting 100,000 DU145 tumor cells suspended in Matrigel into the mice’s hind leg. Once the tumor size reached 100 mm³, the mice were randomized into four groups: Control, RT Only (5x 2Gy irradiation), CB-839 Only (3 x 180mg/kg orally, daily), and CB-839+RT (combination of 3 x 180mg/kg CB-839 orally, daily, and 5x 2Gy irradiation). The solvent for dissolving CB-839 in the study was 25% (w/v) hydroxypropyl-β-cyclodextrin (HPBCD) in 10 mmol/L citrate, pH 2. The dosage was based on previously published studies for various tumor models (56, 57, 59). Tumor size was measured once per week before treatment, using electronic calipers to determine length (L), width (W), and height (H). After treatment commenced, tumor measurements were taken twice weekly. Tumor volume (V) was calculated using the hemiellipsoid equation: V = (L * W * H)/2. The maximal allowed tumor size and burden were not exceeded. At the end of the third day of treatment, blood samples were collected from the mice via retrobulbar blood sampling to isolate plasma for subsequent analysis. Blood was drawn into EDTA-treated collection tubes and centrifuged for 5 minutes at 1,000 rpm. After centrifugation, the plasma was carefully transferred to new tubes and rapidly frozen in liquid nitrogen. The samples were stored at or below −80°C until further processing. Tumor growth was monitored for 200 days or until the tumors reached the maximum allowable volume.

### Statistical Analysis

The results of the flow cytometry analyses, sphere formation assay, Cell cycle assay, γH2A.X assay, Annexin V assay, Mitosox Assay, CM-H2DCFDA assay, Cell-Titer Glo viability assay, Metabolomics data for Parental/Radioresistant cells and CB-839-treated/DMSO-treated datasets, and relative gene expression determined by qPCR were analyzed by paired two-tailed t-test. Statistical analysis for animal experiments was performed using unpaired two-tailed t-test. Outliers were evaluated by using Dixon’s Q test. Additional information about specific statistical analyses is included in the figure legends. The cell survival curves were analyzed using SPSS v.23 software by linear-quadratic formula S(D)/S(0) = exp(-αD-βD2) using stratified linear regression after transformation by the natural logarithm. A significant difference between two survival curves was determined by GraphPad Prism software. A significant difference between the two conditions defined as ns p> 0.05, *p < 0.05; **p < 0.01; ***p < 0.001, ****p < 0.0001. The correlation of gene expression levels was evaluated by SUMO software using the Pearson or Spearman (for nonparametric data) correlation coefficient.

## Results

### GLS activity regulates glutaminolysis and PCa radioresistance *in vivo*

First, we examined whether inhibiting glutaminolysis *in vivo* could enhance the efficacy of RT in murine xenograft models of PCa since the previous *in vitro* studies from our team and others had shown that Gln starvation could radiosensitize PCa cells *in vitro* (21). GLS activity mediates the conversion of Gln to Glu, which is required for GSH production. We hypothesized that GLS inhibition should induce tumor radiosensitization due to increased oxidative stress (Figure 1A). To test our hypothesis, we used CB-839, the only GLS inhibitor that had entered clinical trials and demonstrated promising efficacy and safety in clinical studies (18). To evaluate the radiosensitizing potential of CB-839, we xenografted DU145 cells into the hind leg of immunodeficient Rj:NMRI-*Foxn1^nu/nu^* mice. DU145 cell line was chosen for injection as it has been shown to be dependent on glutaminolysis for the maintenance of the redox state (21). Mice were randomized into four treatment groups: vehicle control, CB-839 alone, RT alone (2 Gy for five consecutive days), and combination therapy (Figure 1B). The combination therapy group exhibited a trend toward prolonged survival compared to the control and other treatment groups (Supplementary Figure 1A). A calculation of tumor volume doubling time and modeling of the tumor volume fold changes revealed statistically significant differences in combination treatment compared to all other groups (Figures 1C, and 1D). Analysis of blood plasma TCA cycle metabolites from the different experimental groups confirmed that Gln levels increased significantly in the CB-839-treated groups (CB-839 alone and combination therapy) compared to control (Supplementary Figure 1B). GLS activity, indicated by the glutamate (Glu)/Gln ratio, was significantly reduced in the CB-839 alone and the combination therapy groups compared to the control, demonstrating systemic inhibition of GLS activity by CB-839 treatment (Figure 1E).

**Figure 1.**
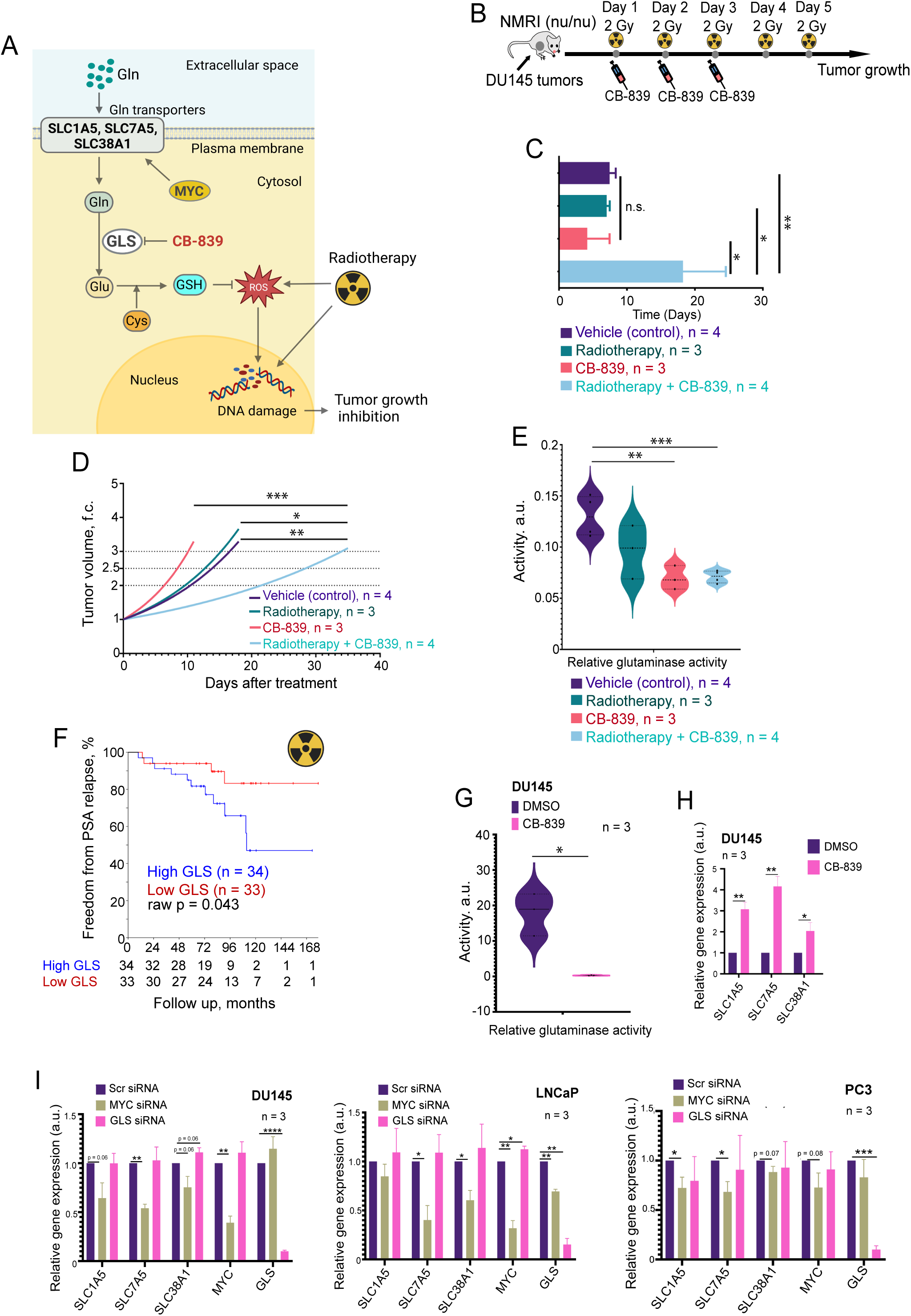
Inhibition of glutaminase activity induces metabolic reprogramming, enhances radiotherapy efficacy and correlates with prostate cancer patient outcomes. **(A)** Schematic representation depicting the role of Gln transporters in the regulation of intracellular Gln homeostasis and mechanism of action of GLS inhibitor, CB-839, on PCa radiosensitization. **(B)** Schematic representation of the *in vivo* experimental design. DU145 prostate cancer cells were subcutaneously implanted into the hind leg of Immunodeficient Rj:NMRI-*Foxn1^nu/nu^* mice. Once tumors reached approximately 100 mm³, mice were randomized into four treatment groups: vehicle control, CB-839 alone (oral administration, 180 mg/kg/day for 3 days), radiotherapy (RT) alone (2 Gy/day for 5 consecutive days), and combination therapy (CB-839 plus RT). **(C)** Tumor volume doubling time presented as a bar chart. Error bars represent standard deviation (SD). The combination therapy group showed a statistically significant increase in tumor doubling time compared to control and other treatment groups (*p < 0.05, **p < 0.01). **(D)** Tumor growth modeling based on measured tumor volumes, depicted as tumor volume fold change over time. The combination therapy significantly delayed tumor growth compared to other groups (*p < 0.05, **p < 0.01, ***p <0.001). **(E)** Glutaminase (GLS) activity calculated as the plasma Glu/Gln ratio. CB-839-treated groups exhibited significantly reduced GLS activity compared to vehicle control group (**p < 0.01, ***p<0.001). **(F)** Kaplan–Meier survival curves for prostate cancer patients (n = 67) with high and low GLS expression levels from the OncoRay cohort. Patients with high GLS expression had significantly higher rates of prostate-specific antigen (PSA) relapse compared to those with low GLS expression (*p < 0.05). **(G)** Violin plots showing glutaminase (GLS) activity (assessed by the Glu/Gln ratio) in DU145 cells treated with CB-839 (GLS inhibitor) or DMSO (control) (*p<0.05). **(H)** qRT-PCR analysis of *SLC1A5, SLC7A5,* and *SLC38A1* expression in DU145 cells treated with CB-839 or DMSO (control). Error bars represent SD (*p < 0.05, **p < 0.01). **(I)** qRT-PCR analysis of *SLC1A5*, *SLC7A5*, *SLC38A1*, *MYC*, and *GLS* expression in DU145 (left), LNCaP (middle), and PC3 (right) cells upon siRNA-

To validate whether GLS expression correlates with tumor radioresistance in PCa patients, we analyzed its expression levels in tumor tissues of patients with intermediate or high-risk localized PCa treated with RT (n = 67). We found a significant association between high GLS expression and a higher risk of PSA relapse, where high-GLS-expressing patients relapsed sooner (Figure 1F). These findings suggest that the level of GLS expression is a marker of PCa sensitivity to radiation therapy and that GLS-driven glutaminolysis is a promising target for PCa radiosensitization.

The cellular uptake of Gln requires amino acid transporters (AATs). We have used a multiparametric analysis to identify clinically relevant AATs playing a role in the regulation of PCa radioresistance (Supplementary Figure 2A-D). We have identified three Gln transporters, *SLC1A5*, *SLC7A5*, and *SLC38A1*, which meet three criteria: (i) their genetic silencing increases radiosensitivity in three analyzed PCa cell lines, (ii) their expression levels significantly correlate with clinical outcome of the patients with PCa, and (iii) their expression levels are upregulated in at least one PCa model with acquired radioresistance (Figure 2A, Supplementary Figure 2A-D, and Supplementary Figure 3AB).

**Figure 2.**
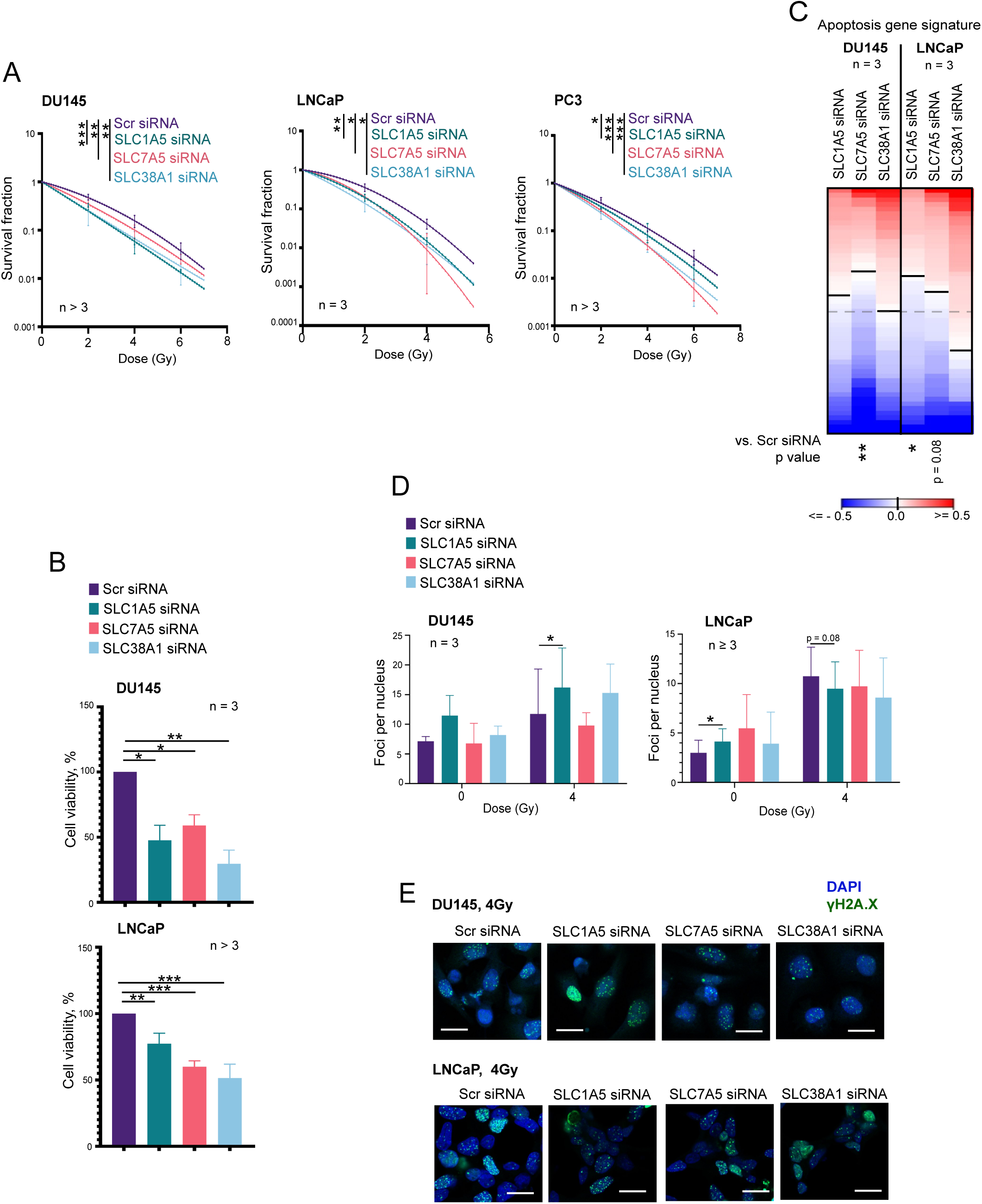
Depletion of amino acid transporters reduces cell viability and enhances radiosensitivity in prostate cancer cells. **(A)** Clonogenic survival curves for DU145 (left), LNCaP (middle), and PC3 (right) cells following transporter knockdown and exposure to increasing doses of radiation (0–6 Gy). Survival fractions were calculated, and curves were fitted using the linear-quadratic model S(D)/S(0) = exp(–αD – βD²). Error bars represent SD (*p < 0.05, **p < 0.01, ***p < 0.001). **(B)** Cell viability of DU145 and LNCaP cells upon siRNA-mediated knockdown of *SLC1A5*, *SLC7A5*, and *SLC38A1*, measured by CellTiter-Glo assay. Data are presented as a percentage normalized to control (siSCR). Error bars represent SD (*p < 0.05, **p < 0.01, ***p < 0.001). **(C)** Analysis of the expression changes of 84 apoptosis-related genes listed in Supplementary Table 10 in response to *SLC1A5, SLC7A5* or *SLC38A1* knockdown compared to Scr siRNA control. **(D)** Bar chart showing the number of γH2AX foci per nucleus in DU145 and LNCaP cells following transporter depletion under sham-irradiated and 4 Gy irradiated conditions. Error bars represent SD (*p < 0.05). **(E)** Representative examples of the γ-H2A.X foci staining in DU145 and LNCaP cells transfected with *SLC1A5*, *SLC7A5*, and *SLC38A1 siRNAs.* Cells transfected with Scr siRNA were used as a control. The images were taken 24 h after cell irradiation with 4 Gy of X-rays. The scale bar is 20 µm.

PCa addiction to Gln is driven by oncogenes, such as *MYC*, which promotes activation of Gln uptake and glutaminolysis (18). The mRNA levels of *SLC1A5*, *SLC7A5*, and *SLC38A1* are upregulated in response to siRNA-mediated MYC knockdown and GLS chemical inhibition (Figure 1G-I). In turn, the genetic silencing of *SLC1A5*, *SLC7A5*, or *SLC38A1* deregulates the expression of *MYC*, *GLS*, and other AATs by a feedback mechanism (Supplementary Figure 4). The analyzed AATs, including *SLC1A5*, *SLC7A5*, or *SLC38A1*, are also upregulated under Gln deprivation in parental PCa cell lines and their derivatives with acquired radioresistance, as discussed in Supplementary Results and as shown in Supplementary Figure 5A-C. These results suggested that the dynamic expression of Gln AATs could be potential tumor survival mechanisms and motivated further exploration of their role in PCa radioresistance.

### Depletion of amino acid transporters reduces viability and enhances radiosensitivity

After establishing that the AATs are upregulated under nutrient stress and in the radioresistant cell models, we aimed to determine if depleting these transporters would affect cell survival and response to RT. The clonogenic survival assays demonstrated that siRNA-mediated targeting of several analyzed Gln AATs significantly sensitized three analyzed cell lines to ionizing radiation (Figure 2A, Supplementary Figure 3). Furthermore, siRNA-mediated knockdown of *SLC1A5*, *SLC7A5*, and *SLC38A1* in DU145 and LNCaP cells significantly reduced cell viability (Figure 2B) and increased the populations of the necrotic and apoptotic cells (Supplementary Figure 6A). We also found significant deregulation of gene signature consisting of 84 genes positively or negatively regulating apoptosis in response to *SLC7A5* or *SLC1A5* knockdown. This observation suggests that the role of Gln transporters in the regulation of PCa cell survival is associated with substantial transcriptomic changes (Figure 2C).

Our previous studies revealed that in HNSCC cell lines, *SLC7A5* regulates radioresistance by activation of the mTORC1 signaling pathway (35). Similar, the knockdown of any of the three transporters, *SLC1A5*, *SLC7A5*, and *SLC38A1,* also led to mTORC1 inhibition in all examined PCa cells (Supplementary Figure 6B). The sensing of amino acid by mTORC1 is one of the key mechanisms regulating cell cycle progression (63). In DU145 cells, *SLC1A5* and *SLC38A1* depletion reduced the G0/G1 phase population, whereas in LNCaP cells, *SLC38A1* depletion slightly reduced the S phase population, as shown in Supplementary Figures 6C and 6D. The cells have been proven to be more radioresistant in G0, early G1, and S phases (64), therefore, cell cycle redistribution by AAT knockdown might contribute to their sensitivity to RT.

Next, we performed the analysis of the γH2AX foci as a marker of DNA double-strand break repair. γH2AX staining revealed that *SLC1A5* depletion led to an increased DNA damage in irradiated DU145 cells (Figures 2D and 2E). In LNCaP cells, *SLC1A5* depletion increased basal DNA damage in sham-irradiated cells, with minimal change post-irradiation (Figures 2D and 2E) suggesting that LNCaP cells depend less on Gln for DNA repair. This observation is consistent with our previously published observations that LNCaP cells can overcome deficient Gln uptake via autophagy (21).

### Transporter depletion reduces cancer stem cell-like properties

In our previous studies, we revealed that Gln deprivation might cause epigenetic reprogramming of PCa cells through the regulation of epigenetic enzymes, such as α-KG-dependent dioxygenases. Many of these enzymes govern the maintenance of cancer stem cell (CSC) populations (21). This is particularly true for PCa cells, such as DU145, whose intracellular pool of α-KG depends on the Gln uptake. In contrast to DU145 cells, LNCaP cells do not rely on the Gln uptake for the α-KG production (21). Given the link between stemness and therapy resistance, we tested whether transporter knockdown also affects CSC-like properties by using sphere formation and Aldefluor assays (Figure 3A). Many analyzed genes encoding the epigenetic enzymes are deregulated after the depletion of AATs, with a greater extent for DU145 cells (Figure 3B). Consistent with our previous finding, the depletion of *SLC1A5*, *SLC7A5*, and *SLC38A1* in DU145 cells significantly reduced spherogenicity in both sham-irradiated and 4Gy irradiated conditions (Figure 3C). These effects align with the inhibition of G1/G0 populations by AAT knockdown, as shown in Supplementary Figures 6C and 6D, since CSC-enriched spherogenic populations in DU145 cells are reported to be quiescent (65). In LNCaP cells, AAT depletion did not induce a statistically significant reduction in spherogenicity, neither in sham-irradiated cells nor in cells irradiated with 4 Gy (Figure 3D). Aldefluor assays revealed that *SLC7A5* and *SLC38A1* depletion decreased the ALDH-positive population in DU145 cells, indicating a reduction in stem-like properties (Figure 3E). In LNCaP cells, there were no statistically significant changes in ALDH-positive populations after AAT knockdown (Figure 3F). Overall, the reduction in stem-like properties upon transporter knockdown highlights the potential of targeting amino acid transporters in cells to eradicate CSC populations. This inhibition is specifically efficient in cells dependent on the Gln uptake for their intracellular α-KG pool.

**Figure 3.**
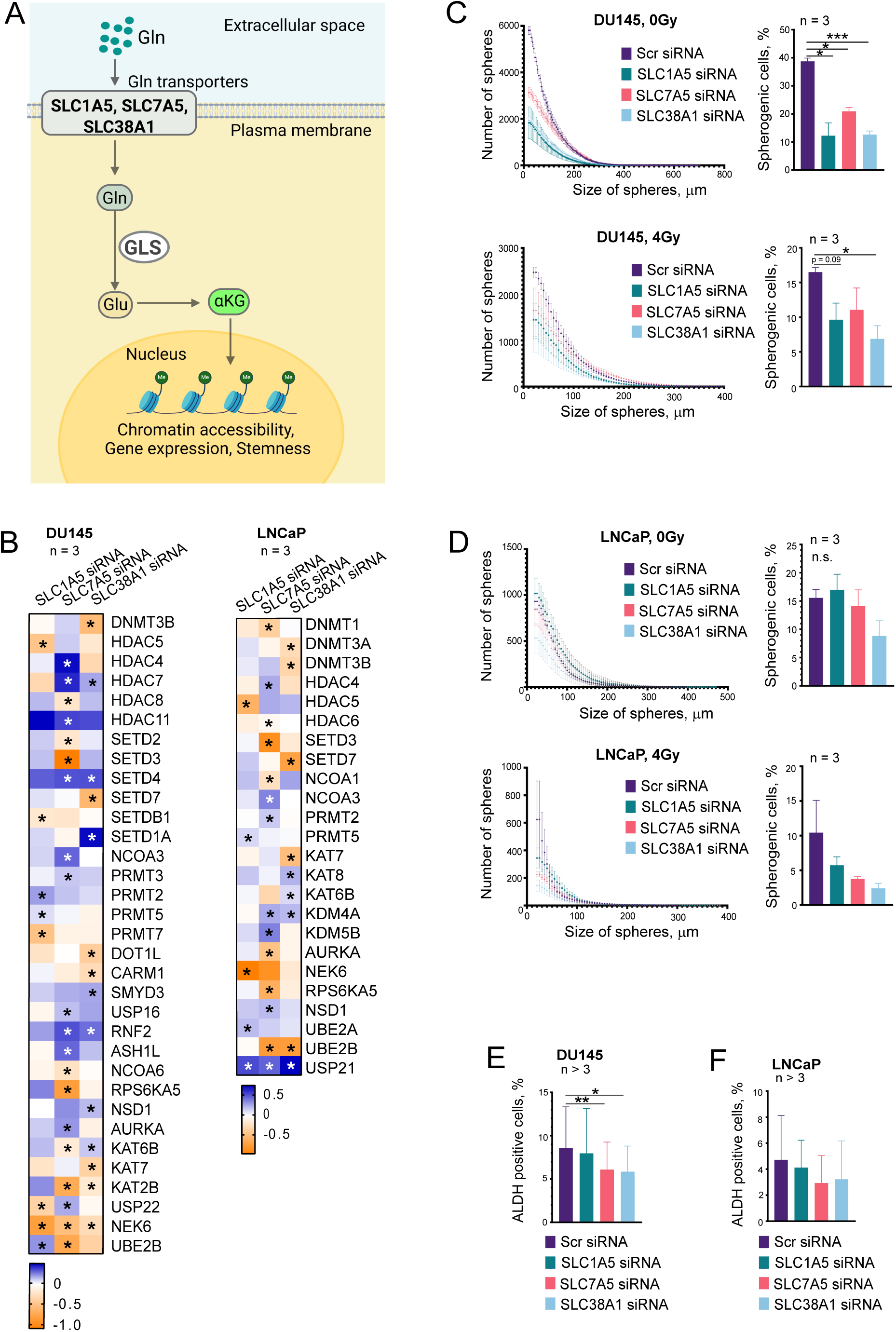
Depletion of amino acid transporters affects stem cell-like properties in PCa cells. **(A)** Schematic representation depicting the role of Gln transporters in the regulation of intracellular αKG levels as a result of GLS-driven catabolism of Gln. **(B)** Expression of epigenetic regulators in response to *SLC1A5, SLC7A5* or *SLC38A1* knockdown compared to Scr siRNA control. Asterisks represent the statistically significant changes. **(C)** Inverse cumulative distribution plots showing sphere size versus number in DU145 cells under sham, and 4Gy irradiation following transporter knockdown. Error bars represent SEM. The bar chart represents the percentage of spherogenic cells capable of forming spheres for each cell line. Error bars represent SD (*p < 0.05, ***p < 0.001). **(D)** Inverse cumulative distribution plots showing sphere size versus number in LNCaP cells under sham, and 4Gy irradiation following transporter knockdown. Error bars represent SEM. The bar chart represents the percentage of spherogenic cells capable of forming spheres for each cell line. Error bars represent SD. **(E)** Aldefluor assay results showing the percentage of ALDH-positive cells in DU145 cells following transporter knockdown. Error bars represent SD (*p < 0.05, **p<0.01). **(F)** Aldefluor assay results showing the percentage of ALDH-positive cells in LNCaP cells following transporter knockdown. Error bars represent SD.

### Targeting Gln transporters induces metabolic reprogramming

Gene expression analyses revealed deregulation of the mitochondrial energy metabolism gene signature after the depletion of AATs (Figure 4A), and Gene set enrichment analysis (GSEA) confirmed upregulation of the oxidative phosphorylation (OXPHOS) genes (Figure 4B), suggesting the adaptive cellular responses to AATs knockdown. Similar to our findings, a previous study revealed an upregulation of mitochondrial respiration in response to amino acid starvation as a compensatory mechanism for the replenishment of protein synthesis (66).

**Figure 4.**
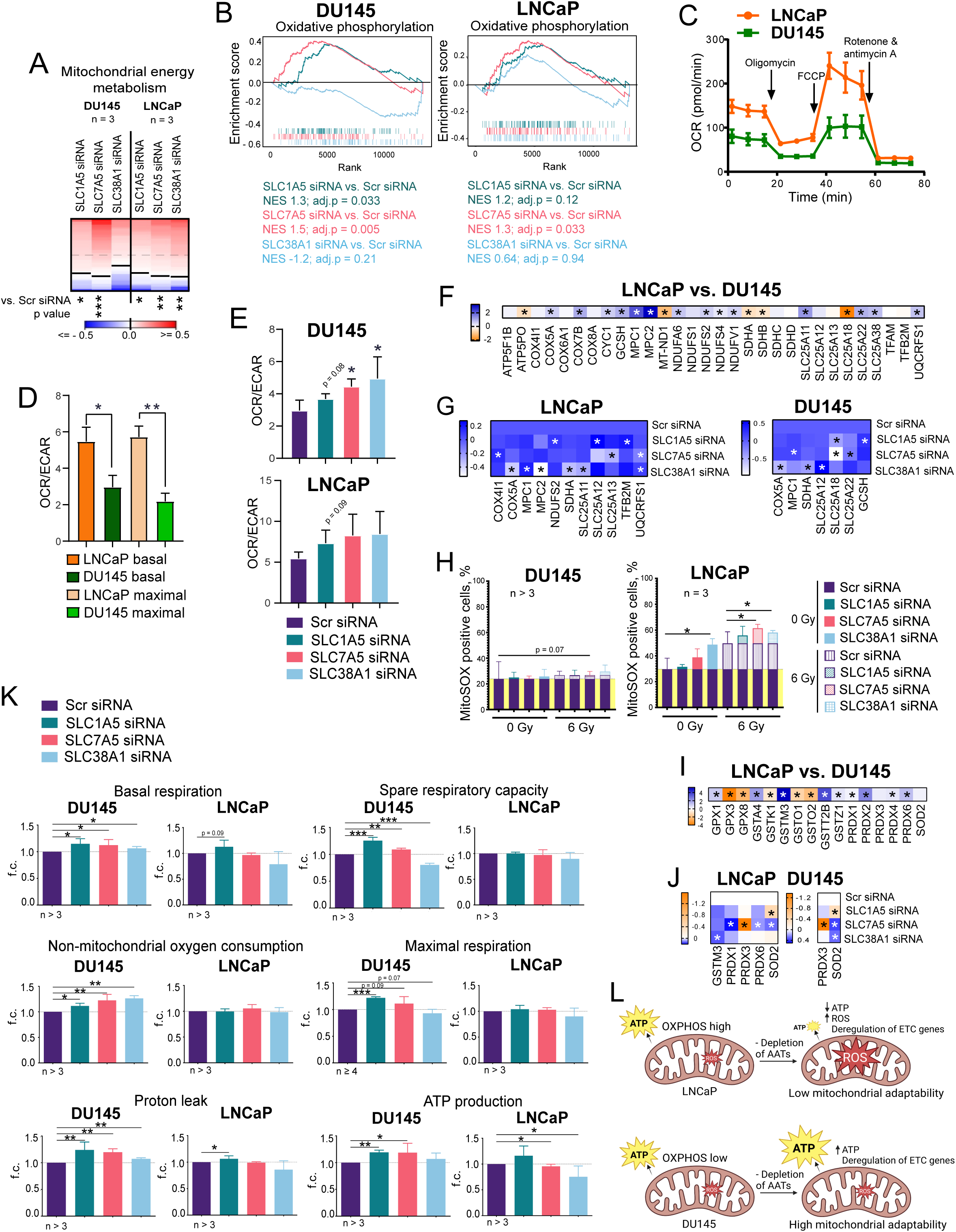
Depletion of amino acid transporters induces metabolic reprogramming. **(A)** Analysis of the expression changes of 79 genes related to the mitochondrial energy metabolism listed in Supplementary Table 10 in response to *SLC1A5, SLC7A5* or *SLC38A1* knockdown compared to Scr siRNA control. **(B)** Gene Set Enrichment Analysis (GSEA) for genes significantly up- or down-regulated upon *SLC1A5*, *SLC7A5*, *SLC38A1* knockdown revealed that deregulated genes are associated with an activation of oxidative phosphorylation. **(C)** The oxygen consumption rate (OCR) was measured using a Seahorse XF Analyzer in DU145 and LNCaP cells. **(D)** The ratio of oxygen consumption rate (OCR) to Extracellular acidification rate (ECAR) in DU145 and LNCaP cells. The OCR/ECAR ratio is a metabolic barometer, quantifying the balance between mitochondrial respiration and glycolysis. **(E)** The ratio of oxygen consumption rate (OCR) to Extracellular acidification rate (ECAR) in DU145 and LNCaP cells in response to *SLC1A5, SLC7A5* or *SLC38A1* knockdown compared to Scr siRNA control. The OCR/ECAR ratio is a metabolic barometer, quantifying the balance between mitochondrial respiration and glycolysis. **(F)** Differential expression of genes playing role in Mitochondrial Energy production and metabolism in LNCaP vs. DU145 cells. **(G)** Effect of the *SLC1A5*, *SLC7A5*, *SLC38A1* knockdown in the expression of genes playing role in Mitochondrial Energy Metabolism in DU145 and LNCaP cells. Asterisks represent the statistically significant changes. **(H)** Mitochondrial superoxide levels in DU145 and LNCaP cells measured by MitoSOX Red staining following transporter knockdown under sham and 6Gy-irradiated conditions. Error bars represent SD (*p < 0.05). **(I)** Differential expression of genes involved in cellular antioxidant defense and redox homeostasis in LNCaP vs. DU145 cells. Asterisks represent the statistically significant changes. **(J)** Effect of the *SLC1A5*, *SLC7A5*, *SLC38A1* knockdown in the expression of genes involved in cellular antioxidant defense and redox homeostasis in LNCaP and DU145 cells. Asterisks represent the statistically significant changes. **(K)** Mitochondrial respiration parameters in DU145 and LNCaP cells following transporter knockdown, measured using the SeaHorse XF MitoStress test kit. Parameters include non-mitochondrial oxygen consumption, ATP production, basal respiration, spare respiratory capacity, maximal respiration, and proton leak. Error bars represent SD (*p < 0.05, **p < 0.01, ***p<0.001). **(L)** Scheme demonstrating the effects of AATs on the mitochondrial adaptability of PCa cells against ROS.

To assess the effects of transporter depletion on cellular metabolism, we used Seahorse XF Analyzer Cell Mito Stress assay. The mitochondrial respiration and glycolysis were analyzed by measuring the oxygen consumption rate (OCR) and extracellular acidification rate (EACR) (Figure 4C, Supplementary Figure 7A). Our previous study and current analyses show that while DU145 cells exhibit a more glycolytic phenotype, LNCaP cells rely on OXPHOS and have high OCR at both basal and maximal mitochondrial respiration states (Figures 4C and 4D, Supplementary Figure 7A). Analysis of the basal bioenergetics state as OCR/ECAR ratios showed increased dependence on OXPHOS after the knockdown of AATs with a higher extent for LNCaP cells (Figure 4E, Supplementary Figure 7B). Most analyzed genes encoding the ETC proteins, mitochondrial amino acid transporters, and mitochondrial transcription factors are upregulated in LNCaP cells (Figure 4F). Many of these genes are significantly deregulated after AAT knockdown, and this deregulation is more pronounced in LNCaP cells (Figure 4G). A comparative phosphoproteomics analysis described in Supplementary Results showed an upregulated phosphorylation of several key mitochondrial proteins in response to AAT depletion in LNCaP and DU145 cells (Supplementary Table 5 and Supplementary Figure 8). These observations suggest that cells with deficient AAT functions activate catabolic processes in mitochondria to maintain cell proliferation, and these compensatory mechanisms are more highly activated in cells with high basal levels of OXPHOS, such as LNCaP.

Previous studies demonstrated chronic oxidative stress and high ROS production in tumor cells with a high OXPHOS phenotype (67). Indeed, LNCaP cells possess a higher level of mitochondria superoxide after depletion of AATs, especially after exposure of cells to RT, a potent inducer of oxidative stress, while DU145 showed no significant changes (Figure 4H). Analysis of the intracellular ROS formation showed the opposite trend with a modest but significant increase in the ROS levels in DU145. In contrast, LNCaP cells showed a highly efficient neutralizing of the intracellular ROS production, which was activated after AATs depletion (Supplementary Figure 7C). These results are in line with our previous observations that LNCaP could compensate for the intracellular level of GSH through the autophagic pathway (21). Expression levels of the antioxidant enzymes were also higher in LNCaP cells and increased in both DU145 and LNCaP cell lines after the knockdown of AATs suggesting an activated response to oxidative stress (Figures 4I and 4J). A significantly increased basal respiration and non-mitochondrial oxygen consumption in DU145 cells, but not in LNCaP after AATs knockdown, also indirectly indicate a high level of non-mitochondrial ROS production (Figure 4K).

Consistent with the deregulation of ETC proteins (Figure 4G), depletion of AATs resulted in a significant increase of proton leak in DU145 after the knockdown of *SLC1A5* and *SLC7A5* and in LNCaP after knockdown of *SLC1A5*. An increase in spare respiratory capacity and maximal respiration in DU145 cells after AAT knockdown suggest a higher mitochondrial adaptation capacity (Figure 4K). Contrary to DU145, LNCaP cells showed no changes in these parameters, suggesting less flexibility in maintaining mitochondrial bioenergetic levels upon scarce amino acid availability. In line with these findings, the ATP production is increased in DU145 after the depletion of AATs (*SLC1A5* and *SLC7A5*), whereas it is decreased in LNCaP cells after the knockdown of AATs (*SLC7A5* and *SLC38A1*) (Figure 4K). This lower mitochondrial adaptability to stress conditions in LNCaP cells can be attributed to their functioning at the maximal capacity prior to stress conditions leading to mitochondrial dysfunction, energy stress, and mitochondrial ROS accumulation upon amino acid deficiency that is even more increased after RT (Figure 4L).

### NUPR1 is a sensor of the glutaminolysis inhibition

Ingenuity Pathway Analysis identified *NUPR1* as one of the top upstream regulators in response to transporter depletion and Gln starvation in DU145 and LNCaP cells (Supplementary Table 6). *NUPR1* functions as an oncogene in different types of tumors, such as breast, thyroid, brain, and pancreatic cancer (68–70). In contrast, *NUPR1* exhibits tumor-suppressive properties in PCa by downregulating key oncogenic pathways, inhibiting tumor cell proliferation and survival, and inducing apoptosis in response to cellular stress (69, 71, 72). NUPR1 expression is known to be induced in response to various cellular stresses (73). Treatment with CB-839 significantly increased *NUPR1* levels in DU145 cells compared to DMSO-treated controls (Figure 5A), similar to the upregulation of AATs (Figure 1H). These findings were confirmed using the cultures of the patient-derived matched tumor and adjacent normal tissues (benign hyperplasia, BPH). These cultures were treated with different concentrations of CB-839, as we described previously (21). Of note, we found more pronounced upregulation of *SLC1A5* and *SLC7A5* in tumor tissues compared to BPH in response to *GLS1* inhibition (Figure 5B).

**Figure 5.**
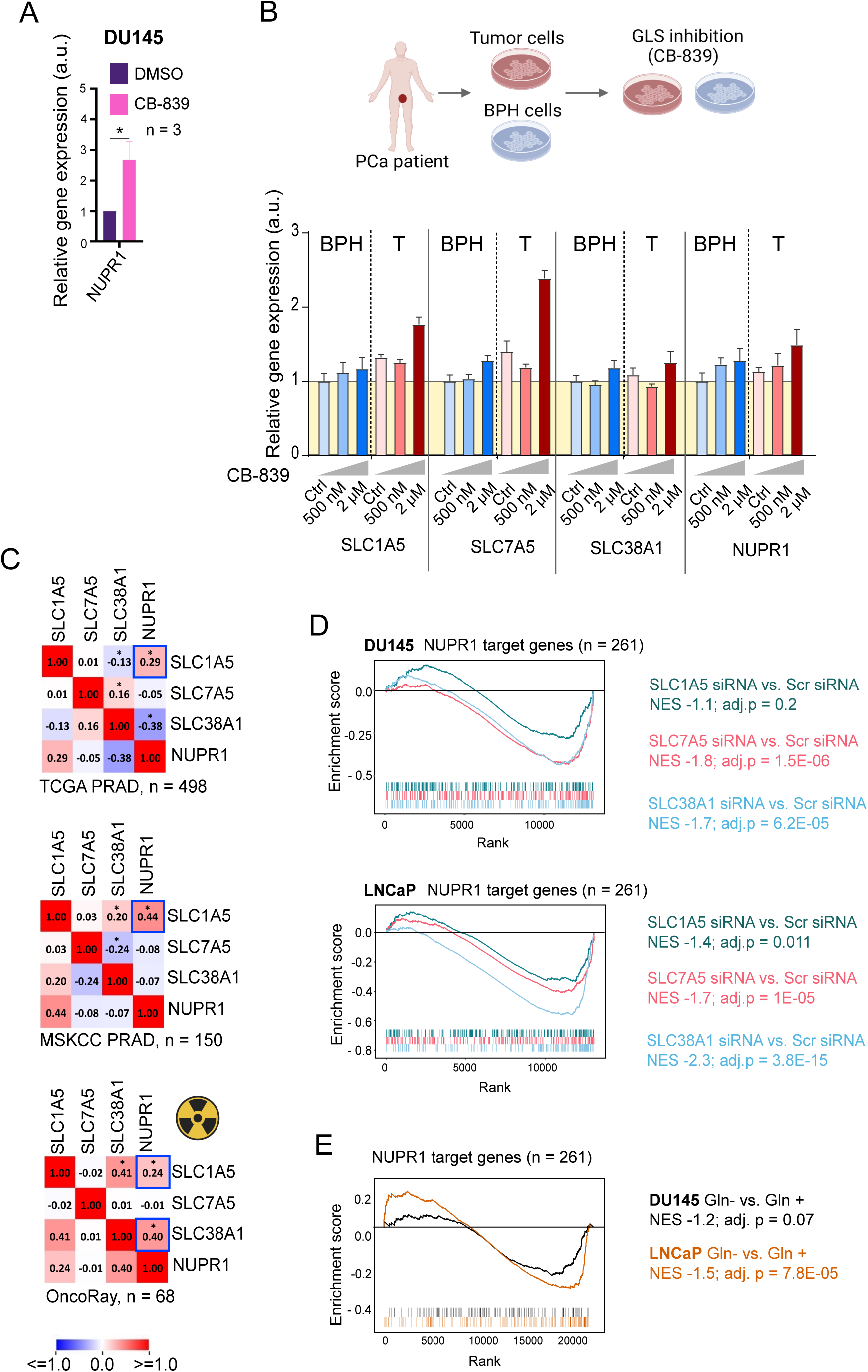
Disrupting Gln metabolism by chemical inhibition of GLS, by Gln starvation or by knockdown of AATs induces NUPR1-mediated stress response, alters the NUPR1-target gene response. **(A)** qRT-PCR analysis of *NUPR1* expression in DU145 cells treated with CB-839 or DMSO (control). Error bars represent SD (*p < 0.05). **(B)** CB-839 treatment of patient-derived tumor (T) tissue (n = 1) and adjacent patient-derived benign prostatic hyperplasia (BPH) (n = 1) increased expression of *SLC1A5, SLC7A5, SLC38A1,* and *NUPR1.* (C) Pearson correlation of mRNA expression levels of *SLC1A5*, *SLC7A5*, *SLC38A1* and *NUPR1* genes in the TCGA PRAD patient cohort (n = 498), MSKCC PRAD patient cohort (n = 150), and OncoRay patient cohort (n = 68), *p < 0.05. **(D)** Gene Set Enrichment Analysis (GSEA) revealed negative enrichment of the NUPR1 target genes (n = 261) in response to *SLC1A5*, *SLC7A5*, and *SLC38A1* knockdown. **(E)** Gene Set Enrichment Analysis (GSEA) revealed negative enrichment of the NUPR1 target genes (n = 261) in response to Gln starvation.

The depletion of *NUPR1* downregulated the expression of all three transporters in LNCaP cells and *SLC38A1* expression in DU145 cells, whereas the knockdown of *SLC7A5* upregulated *NUPR1* expression in DU145 cells, suggesting a feedback regulatory mechanism (Supplementary Figures 9A and 9B). Consistently, we observed a moderate positive correlation of *SLC1A5* and *NUPR1* in the gene expression PCa datasets such as TCGA PRAD (n = 498), MSKCC (n = 150) and OncoRay (n = 68) (Figure 5C).

To further elucidate the molecular pathways linking the *NUPR1* and AATs (*SLC1A5*, *SLC7A5*, and *SLC38A1*), we compiled a master list of *NUPR1* target genes identified from our RNA-seq datasets following transporter depletion in DU145 and LNCaP cells. From the individual gene sets detected by IPA analysis under each condition, we generated a comprehensive list of 261 putative *NUPR1*-regulated genes (Supplementary Table 7). GSEA analysis revealed that this 261-gene subset was negatively enriched in our RNA-seq data for AAT knockdown or Gln depletion (GEO accession numbers GSE288717 and GSE148016 correspondingly), indicating that depletion of transporters or Gln starvation may negatively regulate the *NUPR1*-driven transcriptional program (Figures 5D and 5E). To assess the clinical relevance of this finding, we performed a Pearson correlation analysis using gene expression data from the TCGA PRAD cohort. This analysis identified a core subset of 43 genes whose expression was correlating with all three Gln transporters across patient samples, including 17 negatively correlating genes and 25 positively correlating genes (Supplementary Figure 9C). *NUPR1* is a transcriptional factor whose transcriptional program is activated in response to the nutrient and oxidative stresses (73) (Figure 6A). Gene Ontology (GO) analysis of the 261-gene and 43-gene lists highlighted processes tied to cell cycle regulation, DNA repair, mitosis, nuclear division, and cellular responses to nutrient stress and starvation (Figure 6B, Supplementary Figure 9D, Supplementary Tables 8 and 9). These results demonstrate that *NUPR1* target genes are at an intersection of metabolic and stress response pathways modulated by the depletion of transporters, underlining the importance of *NUPR1* in orchestrating cellular adaptations to metabolic imbalance.

**Figure 6.**
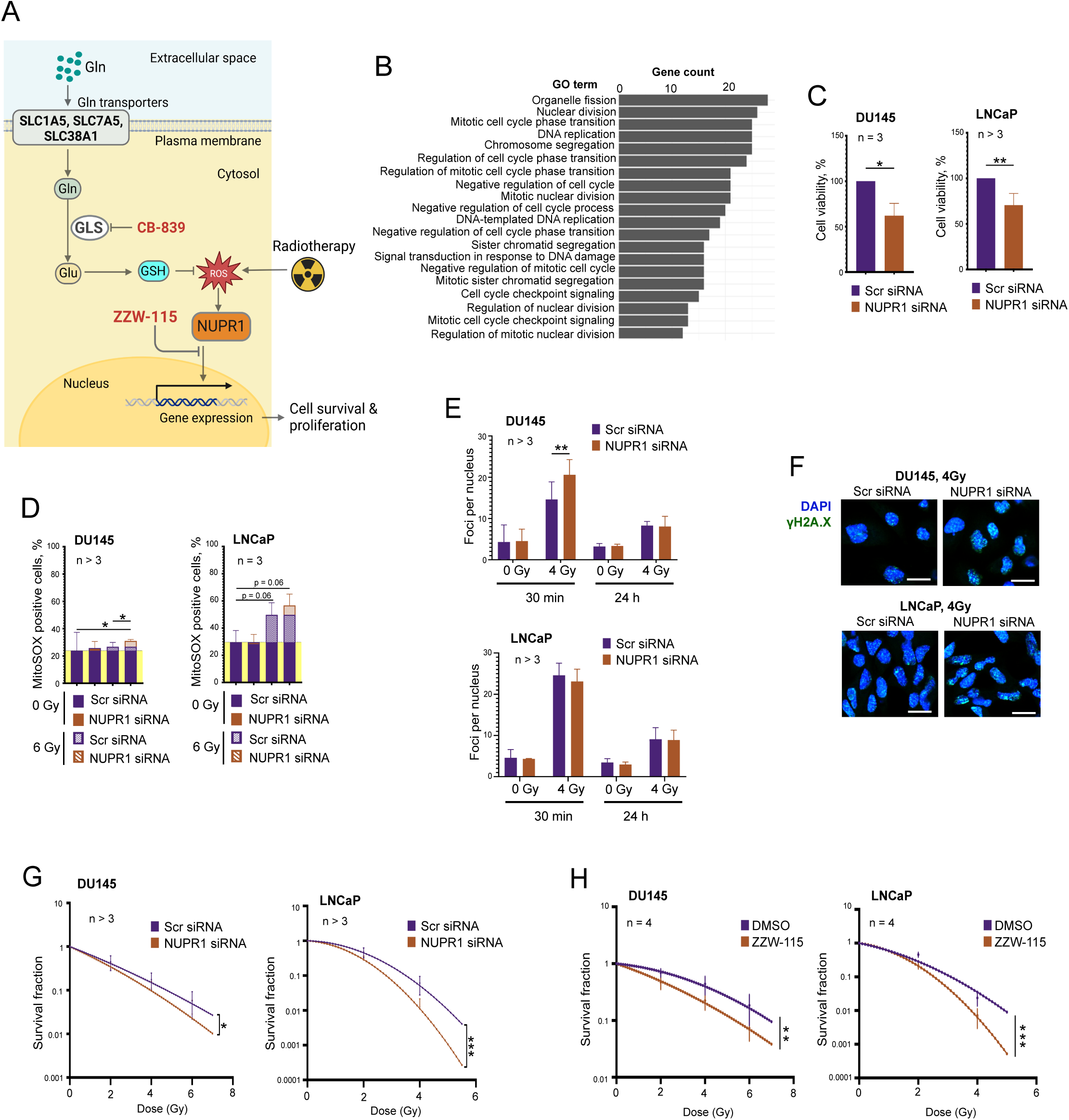
Targeting *NUPR1* reduces cell viability, modulates mitochondrial superoxide levels, and enhances radiosensitivity in prostate cancer cells. **(A)** Schematic representation depicting the role of Gln transporters in the regulation of intracellular Gln homeostasis and mechanism of action of GLS inhibitor, CB-839, and NUPR1 inhibitor, ZZW-115, on PCa radiosensitization. **(B)** A top 20 enriched Gene Ontology (GO) terms in the set of the NUPR1 target genes (n = 261). The gene list is provided in Supplementary Table 7. The statistical data for GO analysis is provided in Supplementary Table 8. **(C)** Cell viability of DU145 and LNCaP cells upon *NUPR1* knockdown, measured by CellTiter-Glo assay. Error bars represent SD (*p < 0.05, **p < 0.01). **(D)** Mitochondrial superoxide levels in DU145 and LNCaP cells measured by MitoSOX Red staining following *NUPR1* knockdown under sham and irradiated conditions. Error bars represent SD (*p < 0.05). **(E)** Bar chart showing the number of γH2AX foci per nucleus in DU145 and LNCaP cells following *NUPR1* depletion under sham-irradiated and 4 Gy irradiated conditions. Error bars represent SD (**p < 0.01). **(F)** Representative examples of the γ-H2A.X foci staining in DU145 and LNCaP cells transfected with *NUPR1 siRNAs.* Cells transfected with Scr siRNA were used as a control. The images were taken 30 min after cell irradiation with 4 Gy of X-rays. The scale bar is 20 µm. **(G)** Clonogenic survival curves for DU145 and LNCaP cells following *NUPR1* knockdown and radiation exposure. Survival fractions were calculated, and curves were fitted using the linear-quadratic model. Error bars represent SD (*p < 0.05, ***p < 0.001). **(H)** Clonogenic survival curves for DU145 and LNCaP cells following ZZW-115 inhibitor treatment and radiation exposure. Survival fractions were calculated, and curves were fitted using the linear-quadratic model. Error bars represent SD (**p < 0.01).

### *NUPR1* depletion reduces viability, alters cell cycle, modulates ROS levels, and enhances radiosensitivity

Given the role of *NUPR1* in stress responses, we explored its functional significance in PCa cell lines. The Cell Titer Glo viability assay showed that *NUPR1* depletion significantly reduced viability and proliferation in both DU145 and LNCaP cells (Figure 6C). Similar to our finding for AATs (Supplementary Figure 6C), cell cycle analysis revealed that *NUPR1* depletion reduced the cell population at the G0/G1 phase and increased the G2/M phase in both sham-irradiated and irradiated DU145 cells (Supplementary Figure 10A). In LNCaP cells, *NUPR1* depletion did not affect G0/G1 phase but slightly increased the S phase population upon irradiation (Supplementary Figure 10B).

MitoSOX Red staining showed that *NUPR1* depletion increased mitochondrial superoxide levels in irradiated DU145 cells. Similar, more pronounced, and close to significant trends were observed In LNCaP cells (Figure 6D). CM-H2DCFDA staining revealed that *NUPR1* depletion decreased ROS levels in LNCaP cells under both sham and irradiated conditions but not in DU145 cells, similar to our observation for AATs (Supplementary Figure 7C and Supplementary Figure 10C).

γH2AX foci analysis indicated that *NUPR1* depletion increased DNA damage in DU145 cells irradiated with 4 Gy 30 minutes after irradiation but not 24 h after irradiation or in LNCaP cells, indicating an elevated initial DNA damage in DU145 cells (Figures 6E and 6F). A radiobiological clonogenic analysis demonstrated that targeting *NUPR1* by using siRNA or the NUPR1 inhibitor, ZZW-115, significantly enhanced radiosensitivity in both PCa cell lines (Figures 6G and 6H, and Supplementary Figures 11A-C).

Analysis of the NUPR1 correlation with RT2 gene sets using the publicly available PCa dataset (PRAD) from The Cancer Genome Atlas (TCGA) (n = 498) (74) revealed the highest correlation with gene signatures related to mitochondrial energy metabolism (Supplementary Figure 11D), suggesting that, similar to the analyzed AATs, *NUPR1* might affect tumor cell viability and sensitivity to radiation by regulation of mitochondrial function.

## Discussion

In our study, we identified amino acid transporters (*SLC1A5*, *SLC7A5*, and *SLC38A1*) and the stress-response gene *NUPR1* as essential regulators of PCa cell survival, metabolic plasticity, and response to RT. By focusing on Gln metabolism, a recognized hallmark of tumor growth and stress adaptation, we uncovered that depletion of these Gln transporters or pharmacological inhibition of GLS disrupts intracellular amino acid homeostasis, compromises redox balance, and ultimately enhances PCa radiosensitivity. Our *in vivo* pilot data further support this concept: targeting Gln catabolism with CB-839, the only clinically approved GLS inhibitor, synergized with RT to delay tumor growth in a PCa xenograft model.

Gln uptake and catabolism are essential for the regulation of key biological functions, such as DNA and protein synthesis, cell proliferation, oxidative stress, mitochondrial homeostasis, and α-KG - driven epigenetic reprogramming (18). Tumor growth is often associated with nutrient limitation due to the high metabolic demands of tumor cells and low nutrient accessibility, and cells might activate different rescue mechanisms to survive the starvation periods (75). The cellular response to the limitation of Gln uptake depends on several factors, including activation of the autophagy as a source of Gln-derived metabolites, dynamic regulation of the AAT as a compensatory mechanism for the replenishment of Gln and other amino acids, and the prevalent energy metabolism (OXPHOS or glycolysis). Our previous and current study demonstrated that PCa cells with activation of ATG-mediated autophagy, such as LNCaP, maintain their intracellular α-KG pool independently of Gln availability (21, 22). These cells demonstrated less effect from the Gln depletion and AAT knockdown at the level of epigenetic reprogramming and CSC regulation in contrast to DU145, whose intracellular availability of α-KG strongly depends on the Gln uptake. Nevertheless, this study revealed a high vulnerability of the cells with high OXPHOS phenotype, such as LNCaP, for the mitochondrial oxidative stress induced by the AAT knockdown, especially after its combination with RT. In line with these results, we previously found significant radiosensitization of LNCaP cells in response to the treatment with metformin, an inhibitor of mitochondrial ETC, independent of the presence of Gln (22). In contrast, DU145 cells exhibit more glycolytic phenotype and possess higher mitochondrial adaptability to the stress conditions induced by AATs. Of importance, both tumor models demonstrated a high plasticity of the AAT expression in response to the knockdown of the single transporter gene and after Gln depletion. Using patient-derived tumor and adjacent non-cancerous tissues, we confirmed that expression of the *SLC1A5*, *SLC7A5*, and *SLC38A1* is a rescue mechanism induced by GLS inhibition. Potentially, plasticity of AAT expression could serve as a marker for cell sensitivity to AAT and GLS inhibition, and tumor cells with less transporter plasticity may be more sensitive to such inhibition. Independent of the cell response to the Gln depletion and knockdown of AATs, both analyzed tumor models became significantly more radiosensitive when Gln transport was disrupted. Our results align with prior studies demonstrating the importance of Gln metabolism in tumor progression and therapy response (19, 21, 30, 31, 43, 58).

Our findings further identified *NUPR1* as a stress sensor that integrates metabolic cues from diminished Gln transport in all analyzed PCa models. Genetic silencing of *NUPR1* reduced cell viability, increased DNA damage, and augmented radiosensitivity, mirroring the impact of AAT depletion. The moderate but consistent correlation between *NUPR1* and several SLC genes in patient cohorts, along with negative enrichment of *NUPR1*-driven target genes in cells lacking specific Gln transporters, reinforces a model in which *NUPR1* helps orchestrate adaptive responses to nutrient and genotoxic stress. Although *NUPR1* has been reported to act as either a tumor promoter or suppressor depending on the cancer context (51–55, 76), our data suggest that it generally supports PCa cell survival under Gln restriction.

All in all, our study suggests that suppression of the tumor Gln metabolism or inhibition of the NUPR1-mediated adaptive stress response in combination with RT could be a promising clinical strategy for the irradiation of tumor cells. Despite these advances, our study has limitations. The *in vivo* experiments were performed in a single xenograft model with a relatively small cohort size, warranting expanded preclinical work to validate and refine dosing schedules for combination therapy. Additionally, although siRNA-mediated gene knockdown is a powerful research technique, it is not clinically relevant. However, the selective pharmacological targeting of *SLC1A5*, *SLC7A5*, and *SLC38A1* is not yet available, and can be clinically feasible only upon the development of selective inhibitors that avoid systemic toxicity and preserve normal tissue function.

In the future, it will be valuable to investigate how Gln transport inhibitors can potentially synergize with emerging therapeutic concepts in PCa. These include immunomodulatory strategies, where metabolic reprogramming may create a more immunogenic tumor microenvironment, targeting OXPHOS with metformin, as well as other targeted agents (e.g., PARP inhibitors) that exploit tumor-specific DNA repair deficits. Elucidating predictive biomarkers, *NUPR1* signaling or gene signatures correlated with Gln transport could refine patient stratification for clinical trials. Further investigation into autophagy and other salvage pathways in Gln-starved PCa cells may identify additional drug targets that block metabolic escape.

In summary, our work underscores the significance of Gln transporters and the *NUPR1*-mediated stress response in PCa cell survival and radioresistance. Disrupting Gln uptake via *SLC1A5*, *SLC7A5*, or *SLC38A1* depletion, or by inhibiting GLS, enhances oxidative stress and curtails DNA damage repair, thereby sensitizing tumor cells to ionizing radiation. These findings provide a rationale for combining Gln-targeted agents, such as CB-839, with RT in prostate cancer and suggest broader therapeutic potential for coupling metabolic interventions with conventional treatment modalities.

## Conclusions

Targeting *SLC1A5*, *SLC7A5*, *SLC38A1*, and *NUPR1*, along with GLS inhibition, disrupts Gln metabolism, increases oxidative stress, and enhances radiosensitivity in prostate cancer emphasizing the critical roles of Gln transporters and *NUPR1* to support cell survival and radioresistance, and offering promising targets to improve the treatment outcomes.

## List of abbreviations

CSC: cancer stem cell
Gln: glutamine
GLS: glutaminase
Glu: glutamate
GSH: glutathione
ER: endoplasmic reticulum
FADH2: flavin adenine dinucleotide
ISR: integrated stress response
mTOR: mechanistic target of rapamycin
NADH: nicotinamide adenine dinucleotide
NUPR1: Nuclear protein 1
OXPHOS: oxidative phosphorylation
PCa: Prostate cancer
RT: radiation therapy
TCA: tricarboxylic acid
UPR: unfolded protein response

## Declarations Ethics declarations

### Ethics approval and consent to participate

Clinical specimens were collected with informed consent from all subjects. The ethical approvals for these retrospective analyses were granted by the local Ethics Committees and Institutional Review Boards of the Faculty of Medicine, Technische Universität Dresden, Germany

### Consent for publication

All authors have agreed to publish this manuscript.

### Availability of data and materials

The publicly available TCGA PRAD (77) and MSKCC (78) datasets were accessed via cBioportal (https://www.cbioportal.org/). RNA-seq datasets for AAT knockdown or Gln depletion are available through GEO data repository (GEO accession numbers GSE288717 and GSE148016 correspondingly). The datasets used and/or analysed during the current study are available from the corresponding author on reasonable request.

### Competing interest: none

Dr. Mechthild Krause received funding for her research projects by IBA (2016), Merck KGaA (2014-2018 for preclinical study; 2018-2020 for clinical study), Medipan GmbH (2014–2018). Dr. Krause, Dr. Linge and Dr. Löck have been involved in an ongoing publicly funded (German Federal Ministry of Education and Research) project with the companies Medipan, Attomol GmbH, GA Generic Assays GmbH, Gesellschaft für medizinische und wissenschaftliche genetische Analysen, Lipotype GmbH and PolyAn GmbH (2019-2021). For the present manuscript, none of the above mentioned funding sources were involved.

### Funding

Work in AD lab was supported by grants from Förderungen des Bundesministeriums für Bildung und Forschung (BMBF) #01DK24011 (COMBOGEGENKREBS), and Deutsche Forschungsgemeinschaft (DFG) #416001651.

### Authors’ contributions

UK and AD conceived the study. UK, VL, IG, MMW, ASK, BA, DS, MP performed the experiments; UK, VL, BA, AL, SL, IIS, MP and AD analyzed the data; IIS, MK and AD supervised the experiments; UK and AD wrote the manuscript; AD supervised the study and provided project administration. All the authors reviewed the final version.

## Acknowledgments

We thank Dr. Kerstin Brüchner for her help on the animal study application. We thank the Light Microscopy Facility of the Center for Molecular and Cellular Bioengineering in Dresden for assistance with the immunofluorescence analyses.

## Notes

### Competing Interest Statement

The authors have declared no competing interest.

## References

1. Bray F, Laversanne M, Sung H, Ferlay J, Siegel RL, Soerjomataram I, et al. Global cancer statistics 2022: GLOBOCAN estimates of incidence and mortality worldwide for 36 cancers in 185 countries. CA Cancer J Clin. 2024;74(3):229–63.

2. Butof R, Dubrovska A, Baumann M. Clinical perspectives of cancer stem cell research in radiation oncology. Radiother Oncol. 2013;108(3):388–96.

3. Schulz A, Meyer F, Dubrovska A, Borgmann K. Cancer Stem Cells and Radioresistance: DNA Repair and Beyond. Cancers (Basel). 2019;11(6).

4. Harris WP, Mostaghel EA, Nelson PS, Montgomery B. Androgen deprivation therapy: progress in understanding mechanisms of resistance and optimizing androgen depletion. Nat Clin Pract Urol. 2009;6(2):76–85.

5. Coller HA. Is cancer a metabolic disease? Am J Pathol. 2014;184(1):4–17.

6. Pavlova NN, Thompson CB. The Emerging Hallmarks of Cancer Metabolism. Cell Metab. 2016;23(1):27–47.

7. Kahya U, Koseer AS, Dubrovska A. Amino Acid Transporters on the Guard of Cell Genome and Epigenome. Cancers (Basel). 2021;13(1).

8. Hanahan D, Weinberg RA. The hallmarks of cancer. Cell. 2000;100(1):57–70.

9. Hanahan D, Weinberg RA. Hallmarks of cancer: the next generation. Cell. 2011;144(5):646–74.

10. Cruzat V, Macedo Rogero M, Noel Keane K, Curi R, Newsholme P. Glutamine: Metabolism and Immune Function, Supplementation and Clinical Translation. Nutrients. 2018;10(11).

11. Bhutia YD, Babu E, Ramachandran S, Ganapathy V. Amino Acid transporters in cancer and their relevance to “glutamine addiction”: novel targets for the design of a new class of anticancer drugs. Cancer Res. 2015;75(9):1782–8.

12. Wise DR, DeBerardinis RJ, Mancuso A, Sayed N, Zhang XY, Pfeiffer HK, et al. Myc regulates a transcriptional program that stimulates mitochondrial glutaminolysis and leads to glutamine addiction. Proc Natl Acad Sci U S A. 2008;105(48):18782–7.

13. Wise DR, Thompson CB. Glutamine addiction: a new therapeutic target in cancer. Trends Biochem Sci. 2010;35(8):427–33.

14. Altman BJ, Stine ZE, Dang CV. From Krebs to clinic: glutamine metabolism to cancer therapy. Nat Rev Cancer. 2016;16(11):749.

15. DeBerardinis RJ, Cheng T. Q’s next: the diverse functions of glutamine in metabolism, cell biology and cancer. Oncogene. 2010;29(3):313–24.

16. Fendt SM, Bell EL, Keibler MA, Olenchock BA, Mayers JR, Wasylenko TM, et al. Reductive glutamine metabolism is a function of the alpha-ketoglutarate to citrate ratio in cells. Nat Commun. 2013;4:2236.

17. Yang L, Venneti S, Nagrath D. Glutaminolysis: A Hallmark of Cancer Metabolism. Annu Rev Biomed Eng. 2017;19:163–94.

18. Erb HHH, Polishchuk N, Stasyk O, Kahya U, Weigel MM, Dubrovska A. Glutamine Metabolism and Prostate Cancer. Cancers (Basel). 2024;16(16).

19. Beier AK, Ebersbach C, Siciliano T, Scholze J, Hofmann J, Honscheid P, et al. Targeting the glutamine metabolism to suppress cell proliferation in mesenchymal docetaxel-resistant prostate cancer. Oncogene. 2024;43(26):2038–50.

20. Lukey MJ, Katt WP, Cerione RA. Targeting amino acid metabolism for cancer therapy. Drug Discov Today. 2017;22(5):796–804.

21. Mukha A, Kahya U, Dubrovska A. Targeting glutamine metabolism and autophagy: the combination for prostate cancer radiosensitization. Autophagy. 2021;17(11):3879–81.

22. Mukha A, Kahya U, Linge A, Chen O, Lock S, Lukiyanchuk V, et al. GLS-driven glutamine catabolism contributes to prostate cancer radiosensitivity by regulating the redox state, stemness and ATG5-mediated autophagy. Theranostics. 2021;11(16):7844–68.

23. Cormerais Y, Vucetic M, Parks SK, Pouyssegur J. Amino Acid Transporters Are a Vital Focal Point in the Control of mTORC1 Signaling and Cancer. Int J Mol Sci. 2020;22(1).

24. Woo Y, Lee HJ, Jung YM, Jung YJ. mTOR-Mediated Antioxidant Activation in Solid Tumor Radioresistance. J Oncol. 2019;2019:5956867.

25. Huo M, Zhang J, Huang W, Wang Y. Interplay Among Metabolism, Epigenetic Modifications, and Gene Expression in Cancer. Front Cell Dev Biol. 2021;9:793428.

26. Koppula P, Zhuang L, Gan B. Cystine transporter SLC7A11/xCT in cancer: ferroptosis, nutrient dependency, and cancer therapy. Protein Cell. 2021;12(8):599–620.

27. White MA, Lin C, Rajapakshe K, Dong J, Shi Y, Tsouko E, et al. Glutamine Transporters Are Targets of Multiple Oncogenic Signaling Pathways in Prostate Cancer. Mol Cancer Res. 2017;15(8):1017–28.

28. Broer A, Fairweather S, Broer S. Disruption of Amino Acid Homeostasis by Novel ASCT2 Inhibitors Involves Multiple Targets. Front Pharmacol. 2018;9:785.

29. Broer A, Gauthier-Coles G, Rahimi F, van Geldermalsen M, Dorsch D, Wegener A, et al. Ablation of the ASCT2 (SLC1A5) gene encoding a neutral amino acid transporter reveals transporter plasticity and redundancy in cancer cells. J Biol Chem. 2019;294(11):4012–26.

30. Broer A, Rahimi F, Broer S. Deletion of Amino Acid Transporter ASCT2 (SLC1A5) Reveals an Essential Role for Transporters SNAT1 (SLC38A1) and SNAT2 (SLC38A2) to Sustain Glutaminolysis in Cancer Cells. J Biol Chem. 2016;291(25):13194–205.

31. Wang Q, Hardie RA, Hoy AJ, van Geldermalsen M, Gao D, Fazli L, et al. Targeting ASCT2-mediated glutamine uptake blocks prostate cancer growth and tumour development. J Pathol. 2015;236(3):278–89.

32. Yoo HC, Park SJ, Nam M, Kang J, Kim K, Yeo JH, et al. A Variant of SLC1A5 Is a Mitochondrial Glutamine Transporter for Metabolic Reprogramming in Cancer Cells. Cell Metab. 2020;31(2):267–83 e12.

33. Zhang Z, Liu R, Shuai Y, Huang Y, Jin R, Wang X, et al. ASCT2 (SLC1A5)-dependent glutamine uptake is involved in the progression of head and neck squamous cell carcinoma. Br J Cancer. 2020;122(1):82–93.

34. Cormerais Y, Giuliano S, LeFloch R, Front B, Durivault J, Tambutte E, et al. Genetic Disruption of the Multifunctional CD98/LAT1 Complex Demonstrates the Key Role of Essential Amino Acid Transport in the Control of mTORC1 and Tumor Growth. Cancer Res. 2016;76(15):4481–92.

35. Digomann D, Kurth I, Tyutyunnykova A, Chen O, Lock S, Gorodetska I, et al. The CD98 Heavy Chain Is a Marker and Regulator of Head and Neck Squamous Cell Carcinoma Radiosensitivity. Clin Cancer Res. 2019;25(10):3152–63.

36. Digomann D, Linge A, Dubrovska A. SLC3A2/CD98hc, autophagy and tumor radioresistance: a link confirmed. Autophagy. 2019;15(10):1850–1.

37. Arndt C, Loureiro LR, Feldmann A, Jureczek J, Bergmann R, Mathe D, et al. UniCAR T cell immunotherapy enables efficient elimination of radioresistant cancer cells. Oncoimmunology. 2020;9(1):1743036.

38. Koseer AS, Loureiro LR, Jureczek J, Mitwasi N, Gonzalez Soto KE, Aepler J, et al. Validation of CD98hc as a Therapeutic Target for a Combination of Radiation and Immunotherapies in Head and Neck Squamous Cell Carcinoma. Cancers (Basel). 2022;14(7).

39. Schulte ML, Fu A, Zhao P, Li J, Geng L, Smith ST, et al. Pharmacological blockade of ASCT2-dependent glutamine transport leads to antitumor efficacy in preclinical models. Nat Med. 2018;24(2):194–202.

40. Kanai Y, Hediger MA. The glutamate/neutral amino acid transporter family SLC1: molecular, physiological and pharmacological aspects. Pflugers Arch. 2004;447(5):469–79.

41. Nicklin P, Bergman P, Zhang B, Triantafellow E, Wang H, Nyfeler B, et al. Bidirectional transport of amino acids regulates mTOR and autophagy. Cell. 2009;136(3):521–34.

42. Verrey F, Closs EI, Wagner CA, Palacin M, Endou H, Kanai Y. CATs and HATs: the SLC7 family of amino acid transporters. Pflugers Arch. 2004;447(5):532–42.

43. Wang Q, Tiffen J, Bailey CG, Lehman ML, Ritchie W, Fazli L, et al. Targeting amino acid transport in metastatic castration-resistant prostate cancer: effects on cell cycle, cell growth, and tumor development. J Natl Cancer Inst. 2013;105(19):1463–73.

44. Chen R, Zou Y, Mao D, Sun D, Gao G, Shi J, et al. The general amino acid control pathway regulates mTOR and autophagy during serum/glutamine starvation. J Cell Biol. 2014;206(2):173–82.

45. Broer S. The SLC38 family of sodium-amino acid co-transporters. Pflugers Arch. 2014;466(1):155–72.

46. Gao P, Tchernyshyov I, Chang TC, Lee YS, Kita K, Ochi T, et al. c-Myc suppression of miR-23a/b enhances mitochondrial glutaminase expression and glutamine metabolism. Nature. 2009;458(7239):762–5.

47. Dang CV. MYC, metabolism, cell growth, and tumorigenesis. Cold Spring Harb Perspect Med. 2013;3(8).

48. Yue M, Jiang J, Gao P, Liu H, Qing G. Oncogenic MYC Activates a Feedforward Regulatory Loop Promoting Essential Amino Acid Metabolism and Tumorigenesis. Cell Rep. 2017;21(13):3819–32.

49. Han S, Zhu L, Zhu Y, Meng Y, Li J, Song P, et al. Targeting ATF4-dependent pro-survival autophagy to synergize glutaminolysis inhibition. Theranostics. 2021;11(17):8464–79.

50. Jin HO, Hong SE, Kim JY, Jang SK, Park IC. Amino acid deprivation induces AKT activation by inducing GCN2/ATF4/REDD1 axis. Cell Death Dis. 2021;12(12):1127.

51. Gironella M, Malicet C, Cano C, Sandi MJ, Hamidi T, Tauil RM, et al. p8/nupr1 regulates DNA-repair activity after double-strand gamma irradiation-induced DNA damage. J Cell Physiol. 2009;221(3):594–602.

52. He W, Cheng F, Zheng B, Wang J, Zhao G, Yao Z, et al. NUPR1 is a novel potential biomarker and confers resistance to sorafenib in clear cell renal cell carcinoma by increasing stemness and targeting the PTEN/AKT/mTOR pathway. Aging (Albany NY). 2021;13(10):14015–38.

53. Huang C, Santofimia-Castano P, Iovanna J. NUPR1: A Critical Regulator of the Antioxidant System. Cancers (Basel). 2021;13(15).

54. Kim YJ, Hong SE, Jang SK, Park KS, Kim CH, Park IC, et al. Knockdown of NUPR1 Enhances the Sensitivity of Non-small-cell Lung Cancer Cells to Metformin by AKT Inhibition. Anticancer Res. 2022;42(7):3475–81.

55. Zhan Y, Zhang Z, Liu Y, Fang Y, Xie Y, Zheng Y, et al. NUPR1 contributes to radiation resistance by maintaining ROS homeostasis via AhR/CYP signal axis in hepatocellular carcinoma. BMC Med. 2022;20(1):365.

56. Boysen G, Jamshidi-Parsian A, Davis MA, Siegel ER, Simecka CM, Kore RA, et al. Glutaminase inhibitor CB-839 increases radiation sensitivity of lung tumor cells and human lung tumor xenografts in mice. Int J Radiat Biol. 2019;95(4):436–42.

57. Gross MI, Demo SD, Dennison JB, Chen L, Chernov-Rogan T, Goyal B, et al. Antitumor activity of the glutaminase inhibitor CB-839 in triple-negative breast cancer. Mol Cancer Ther. 2014;13(4):890–901.

58. Lampa M, Arlt H, He T, Ospina B, Reeves J, Zhang B, et al. Glutaminase is essential for the growth of triple-negative breast cancer cells with a deregulated glutamine metabolism pathway and its suppression synergizes with mTOR inhibition. PLoS One. 2017;12(9):e0185092.

59. Wicker CA, Hunt BG, Krishnan S, Aziz K, Parajuli S, Palackdharry S, et al. Glutaminase inhibition with telaglenastat (CB-839) improves treatment response in combination with ionizing radiation in head and neck squamous cell carcinoma models. Cancer Lett. 2021;502:180–8.

60. Kurth I, Hein L, Mabert K, Peitzsch C, Koi L, Cojoc M, et al. Cancer stem cell related markers of radioresistance in head and neck squamous cell carcinoma. Oncotarget. 2015;6(33):34494–509.

61. Schindelin J, Arganda-Carreras I, Frise E, Kaynig V, Longair M, Pietzsch T, et al. Fiji: an open-source platform for biological-image analysis. Nat Methods. 2012;9(7):676–82.

62. Gebhardt M, Kunath C, Frobel D, Funk AM, Peitzsch M, Nolting S, et al. Identification of Succinate Dehydrogenase Gene Variant Carriers by Blood Biomarkers. J Endocr Soc. 2024;8(9):bvae142.

63. Rong Y, Darnell AM, Sapp KM, Vander Heiden MG, Spencer SL. Cells use multiple mechanisms for cell-cycle arrest upon withdrawal of individual amino acids. Cell Rep. 2023;42(12):113539.

64. Pawlik TM, Keyomarsi K. Role of cell cycle in mediating sensitivity to radiotherapy. Int J Radiat Oncol Biol Phys. 2004;59(4):928–42.

65. Li S, Goncalves KA, Lyu B, Yuan L, Hu GF. Chemosensitization of prostate cancer stem cells in mice by angiogenin and plexin-B2 inhibitors. Commun Biol. 2020;3(1):26.

66. Johnson MA, Vidoni S, Durigon R, Pearce SF, Rorbach J, He J, et al. Amino acid starvation has opposite effects on mitochondrial and cytosolic protein synthesis. PLoS One. 2014;9(4):e93597.

67. Gentric G, Kieffer Y, Mieulet V, Goundiam O, Bonneau C, Nemati F, et al. PML-Regulated Mitochondrial Metabolism Enhances Chemosensitivity in Human Ovarian Cancers. Cell Metab. 2019;29(1):156–73 e10.

68. Chowdhury UR, Samant RS, Fodstad O, Shevde LA. Emerging role of nuclear protein 1 (NUPR1) in cancer biology. Cancer Metastasis Rev. 2009;28(1-2):225–32.

69. Martin TA, Li AX, Sanders AJ, Ye L, Frewer K, Hargest R, et al. NUPR1 and its potential role in cancer and pathological conditions (Review). Int J Oncol. 2021;58(5).

70. Santofimia-Castano P, Xia Y, Peng L, Velazquez-Campoy A, Abian O, Lan W, et al. Targeting the Stress-Induced Protein NUPR1 to Treat Pancreatic Adenocarcinoma. Cells. 2019;8(11).

71. Jiang WG, Davies G, Martin TA, Kynaston H, Mason MD, Fodstad O. Com-1/p8 acts as a putative tumour suppressor in prostate cancer. Int J Mol Med. 2006;18(5):981–6.

72. Schnepp PM, Shelley G, Dai J, Wakim N, Jiang H, Mizokami A, et al. Single-Cell Transcriptomics Analysis Identifies Nuclear Protein 1 as a Regulator of Docetaxel Resistance in Prostate Cancer Cells. Mol Cancer Res. 2020;18(9):1290–301.

73. Liu S, Costa M. The role of NUPR1 in response to stress and cancer development. Toxicol Appl Pharmacol. 2022;454:116244.

74. Cancer Genome Atlas Research N. The Molecular Taxonomy of Primary Prostate Cancer. Cell. 2015;163(4):1011–25.

75. Pavlova NN, Zhu J, Thompson CB. The hallmarks of cancer metabolism: Still emerging. Cell Metab. 2022;34(3):355–77.

76. Clark DW, Mitra A, Fillmore RA, Jiang WG, Samant RS, Fodstad O, et al. NUPR1 interacts with p53, transcriptionally regulates p21 and rescues breast epithelial cells from doxorubicin-induced genotoxic stress. Curr Cancer Drug Targets. 2008;8(5):421–30.

77. Sanchez-Vega F, Mina M, Armenia J, Chatila WK, Luna A, La KC, et al. Oncogenic Signaling Pathways in The Cancer Genome Atlas. Cell. 2018;173(2):321–37 e10.

78. Taylor BS, Schultz N, Hieronymus H, Gopalan A, Xiao Y, Carver BS, et al. Integrative genomic profiling of human prostate cancer. Cancer Cell. 2010;18(1):11–22.

