## Supplementary material for "Glutamine transporters regulate prostate cancer radiosensitivity through NUPR1-mediated stress response": Figure S 1-11, Table S 1-6

#### Supplementary results

##### Gln starvation and Gln transporter expression in parental (P) and radioresistant (RR) sublines

To clarify whether nutrient deprivation alone drives transporter upregulation, we apply Gln starvation on both P and RR sublines and measured the expression of AATs. In DU145, Gln deprivation significantly increased the expression of *SLC1A5*, *SLC7A5*, *SLC38A1*, and *SLC38A5* in both P and RR cells (Supplementary Figure 5A). Similarly, LNCaP cells, demonstrated increased expression of *SLC1A5*, *SLC7A5*, *SLC38A1*, *SLC38A7*, and *SLC7A7* with more pronounced increases in the RR subline (Supplementary Figure 5B). Additionally, the results of gene expression microarray confirmed significant upregulation of these transporters under Gln starvation (Supplementary Figure 5C). Overall, these findings suggest the activation of adaptive responses to nutrient deprivation.

##### Phosphoproteomics analysis reveals cell line–specific mechanisms of transporter depletion

To elucidate the molecular mechanisms underlying the effects of transporter depletion, we performed phosphoproteomics analysis on DU145 and LNCaP cells with shRNA-mediated knockdown of *SLC1A5*, *SLC7A5*, and *SLC38A1* (Supplementray Table 5). For each gene of interest, two different shRNAs were used, with the non-specific shRNA plasmid as a control. The cells were transfected with the plasmids listed in Supplementary Table 2 and then selected using puromycin to eliminate untransfected and transiently transfected cells. Knockdown efficiency was confirmed by Western blotting (Supplementary Figure 8A) and qPCR (Supplementary Figure 8B). The phosphoproteomic profiling was performed by Omics Technologies Inc. (USA).

We found that transporter depletion in DU145 cells caused significant alterations in the phosphorylation of the proteins associated with mitochondrial function and apoptosis (Supplementary Table 5). Notably, increased TOMM20 and AIFM1 phosphorylation recapitulate impaired mitochondrial protein import and induction of caspase-independent apoptosis, respectively [1-5]. TOMM20, Translocase of the outer mitochondrial membrane 20, is essential to import nuclear-encoded proteins into mitochondria [2]. Increased phosphorylation of TOMM20 can alter the efficiency of mitochondrial protein import, leading to mitochondrial dysfunction. Apoptosis-inducing factor, mitochondrion-associated 1 (AIFM1) is involved in caspase-independent apoptosis by inducing chromatin condensation and DNA fragmentation. Increased phosphorylation of AIFM1 may activate caspase-independent apoptosis pathway linked to mitochondrial dysfunction [1-5]. Also, decreased phosphorylation of HSP90AA1 and SIRT1 indicates compromised mitochondrial protein folding and regulation, leading to mitochondrial stress and dysfunction [6, 7]. These alterations are in line with increased basal respiration and proton leak observed in Seahorse assays (Figure 4K).

On the other hand, LNCaP cells place less emphasis on mitochondrial proteins and more on transcription factors and cell cycle regulators (Supplementary Table 5). LNCaP cells bearing transporter knockdown demonstrated augmented phosphorylation of proteins involved in cell cycle regulation and stress responses, such as FOXM1 and HSF1. FOXM1 is a transcription factor critical for cell cycle progression, DNA repair, and genomic stability [8, 9]. Phosphorylation activates FOXM1, promoting the transcription of genes playing a role in G2/M transition and DNA repair. Enhanced FOXM1 activity can lead to cell cycle dysregulation [8, 9]. Phosphorylation activates HSF1, inducing the expression of heat shock proteins that are important for protein folding and protecting against stress [10]. Enhanced activity of these proteins can alter cell cycle progression and upregulation of heat shock proteins, supporting cell survival under stress [8-10]. In addition, knockdown of each of these AATs in LNCaP cells resulted in the upregulation of phosphorylation of the Inner Membrane Mitochondrial Protein (IMMT), which regulates mitochondrial architecture and function [11, 12], HSF1, which is required for the activation of mitochondrial chaperone genes [13], and FOXM1, a recently described OXPHOS regulator [14, 15].

#### Supplementary material

**Supplementary Tables:**

**Supplementary Table 1.** The siRNA sequences used in the study.

**Supplementary Table 2.** The shRNA expression vectors used in the study.

**Supplementary Table 3.** The primers used in the study.

**Supplementary Table 4.** Antibodies used for Western blot analysis.

**Supplementary Table 5.** A comparative phosphoproteomics analysis of PCa cell lines DU145 and LNCaP with or without knockdown of Gln AATs.

**Supplementary Table 6:** The upstream regulators identified in PCa cell lines DU145 and LNCaP using Ingenuity Pathway Analysis (IPA) in response to specified treatments.

**Supplementary Table 7.** A list of 261 NUPR1 target genes identified using Ingenuity Pathway Analysis (IPA) in the RNA seq datasets for SLC knockdown in DU145 and LNCaP cells.

**Supplementary Table 8.** Gene Ontology (GO) analysis of the 261-gene list of the NUPR1 transcriptional targets.

**Supplementary Table 9.** Gene Ontology (GO) analysis of the 43-gene list of the NUPR1 transcriptional targets

**Supplementary Table 10.** Supplementary Table 6. Gene sets used for the correlative analysis.

**Supplementary figures:**

**Supplementary Figure 1. Inhibition of GLS activity reprograms cell metabolism and inhibits tumor growth. (A)** Kaplan–Meier survival curves showing the probability of survival for each treatment group over time (days after the start of treatment). The combination therapy group exhibited a trend toward prolonged survival compared to other groups. **(B)** Mouse plasma glutamine (Gln) and glutamate (Glu) levels upon different treatment arms. CB-839-treated groups showed significantly increased Gln levels compared to control (* $p < 0.05$).

**Supplementary Figure 2. Overview of multiparametric *in silico* approaches to identify clinically relevant glutamine transporters in prostate cancer. (A)** A brief summary of bioinformatics and computational analyses used to select studied amino acid transporters mediating glutamine transport. The approach included evaluating expression levels, functional assays, and clinical relevance in prostate cancer. **(B)** Radiobiological clonogenic survival curves for DU145 and LNCaP, radioresistant (RR) and parental (P) cell sublines cells that are used in the study. Survival fractions were calculated, and curves were fitted using the linear-quadratic model. Error bars represent SD *** $p < 0.001$). **(C)** Correlation of the expression levels of selected Gln transporters with clinical outcomes in the PCa TCGA cohort (n = 498). OS: overall survival; DFS: disease-free survival: BRFS: biochemical recurrence-free survival. **(D)** Heatmap of microarray data showing transporter expression in DU145 and LNCaP P and RR cells. Red color indicates high expression; blue color indicates low expression.

**Supplementary Figure 3. Plating efficiency and clonogenic survival in prostate cancer cells upon transporter depletion. (A)** Clonogenic survival curves for DU145, LNCaP and PC3 cells upon depletion of indicated transporters and exposure to increasing doses of radiation. Survival fractions were calculated, and curves were fitted using the linear-quadratic model $S(D)/S(0) = \exp(-\alpha D - \beta D^2)$ using stratified linear regression after natural logarithm transformation. Error bars represent SD. (* $p < 0.05$,

**$p<0.01$, ***$<0.001$). **(B)** Bar chart showing plating efficiency (%) of DU145, LNCaP and PC3 cells upon transporter depletion. Error bars represent SD. (*$p < 0.05$).

**Supplementary Figure 4. qRT-PCR analysis of expression of studied transporter and metabolic genes in DU145, LNCaP and PC3 cells upon depletion of *SLC1A5*, *SLC7A5*, and *SLC38A1*.** Error bars represent SD (*$p < 0.05$, **$p<0.01$, ***$<0.001$, ****$<0.0001$).

**Supplementary Figure 5. Differential expression of AATs in response to Gln starvation. (A)** qRT-PCR analysis of transporter expression in DU145 P and RR cells under glutamine starvation (–Gln) and non-starvation (+Gln) conditions. Expression levels are demonstated as fold change relative to +Gln parental cells. Error bars represent SD (*$p < 0.05$, ****$p < 0.0001$). **(B)** qRT-PCR analysis of transporter expression in LNCaP P and RR cells under glutamine starvation (–Gln) and non-starvation (+Gln) conditions. Expression levels are demonstrated as fold change relative to +Gln parental cells. Error bars represent SD (*$p < 0.05$, ***$p < 0.001$, ****$p < 0.0001$). **(C)** Heatmap of microarray gene expression data showing expression of glutamine transporters in DU145 (up) and LNCaP (down) cells under glutamine starvation (–Gln) and non-starvation (+Gln) conditions.

**Supplementary Figure 6. Transporter knockdown alters cell cycle distribution and mTORC1 signaling in PCa cells. (A)** Cell death analysis in DU145 and LNCaP cells using Annexin V/PI staining following transporter knockdown under sham and irradiated (6 Gy) conditions. Bar charts represent percentages of apoptotic and necrotic cells. Error bars represent SD (*$p < 0.05$). **(B)** Gene Set Enrichment Analysis (GSEA) for genes significantly up- or down regulated upon *SLC1A5*, *SLC7A5*, *SLC38A1* knockdown revealed that deregulated genes are associated with an inhibition of mTORC1 signaling. **(C)** Cell cycle distribution of DU145 and LNCaP cells under sham-irradiated conditions following transporter knockdown, analyzed by flow cytometry. Error bars represent SD

(*$p < 0.05$). **(D)** Cell cycle distribution of DU145 and LNCaP cells under 6Gy-irradiated conditions following transporter knockdown, analyzed by flow cytometry. Error bars represent SD (*$p < 0.05$).

**Supplementary Figure 7. Targeting AATs alter intracellular ROS and mitochondrial superoxide levels. (A)** The Extracellular acidification rate (ECAR) was measured using a Seahorse XF Analyzer in DU145 and LNCaP cells. **(B)** The ratio of basal oxygen consumption rate (OCR) to basal ECAR in DU145 and LNCaP cells in response to knockdown of *SLC1A5*, *SLC7A5*, and *SLC38A1*. The OCR/ECAR ratio is a metabolic barometer, quantifying the balance between mitochondrial respiration and glycolysis. **(C)** Reactive oxygen species (ROS) levels in DU145 and LNCaP cells measured by CM-H2DCFDA staining following transporter knockdown under sham (0 Gy) and irradiated (6 Gy) conditions. Error bars represent SD (*$p < 0.05$, **$p<0.01$).

**Supplementary Figure 8. Validation of stable shRNA-mediated knockdown of glutamine transporters in DU145 and LNCaP cells. (A)** Western blot analysis confirming the stable knockdown of SLC1A5, SLC7A5, and SLC38A1 in DU145 cells and LNCaP selected with 5 µg/mL puromycin. GAPDH was used as a loading control. **(B)** qRT-PCR analysis confirming the stable knockdown of *SLC1A5*, *SLC7A5*, and *SLC38A1* in DU145 and LNCaP cells selected with 5 µg/mL puromycin. Expression levels are normalized to *ACTB* and *RPLP0*, and presented as fold change relative to cells transfected with control shRNA vector.

**Supplementary Figure 9. The interplay between SLC genes and NUPR1 expression and transcriptional activity in prostate cancer cells. (A)** qRT-PCR analysis of *SLC1A5*, *SLC7A5*, *SLC38A1*, and *NUPR1* expression in DU145 and LNCaP cells upon siRNA-mediated knockdown of *NUPR1*. Error bars represent SD (*$p < 0.05$, **$p < 0.01$, ****$p < 0.0001$). **(B)** qRT-PCR analysis of *NUPR1* expression in DU145 and LNCaP cells upon siRNA-mediated knockdown of *SLC1A5*, *SLC7A5*, and *SLC38A1*. Error bars represent SD (*$p < 0.05$). **(C)** Pearson correlation of mRNA expression levels of *SLC1A5*, *SLC7A5*, and *SLC38A1* genes in the TCGA PRAD patient cohort (n = 498). **(D)** A top 20 enriched Gene Ontology (GO) terms in the set of the NUPR1 target genes (n = 43). The statistical data for GO analysis is provided in Supplementary Table 9.

**Supplementary Figure 10. Depletion of *NUPR1* alters cell cycle progression, and ROS levels. (A)** Cell cycle distribution of DU145 cells under sham-irradiated conditions or after irradiation with 6 Gy following siRNA-mediated knockdown of *NUPR1*. Error bars represent SD (*$p < 0.05$, **$p < 0.01$). **(B)** Cell cycle distribution of LNCaP cells under sham-irradiated conditions or after irradiation with 6 Gy following siRNA-mediated knockdown of *NUPR1*. Error bars represent SD (*$p < 0.05$). **(C)** ROS levels in DU145 and LNCaP cells measured by CM-H2DCFDA staining following *NUPR1* knockdown under sham and irradiated conditions. Error bars represent SD (*$p<0.05$, **$p < 0.01$).

**Supplementary Figure 11. Effects of *NUPR1* depletion on cancer stem cell properties, cell death, and DNA damage in prostate cancer cells. (A)** Bar chart showing plating efficiency (%) of DU145 and LNCaP cells upon *NUPR1* depletion and ZZW-115 treatment. **(B)** DU145, and LNCaP were treated for 48 h with different doses of *NUPR1* inhibitor ZZW-115 at concentrations ranging from 100 µM to 0 µM and viability calculated via Cell-Titer Glo assay. $IC_{50}$ values (50% lethal dose) were determined for each cell line individually by non-linear regression analysis using GraphPad Prism software (San Diego, USA). **(C)** Bar chart showing plating efficiency (%) of DU145 and LNCaP cells upon ZZW-115 treatment. **(D)** Analysis of the *NUPR1* correlation with RT2 gene sets using the publicly available PCa dataset (PRAD) from The Cancer Genome Atlas (TCGA) (n = 498).

Supplementary Figure 1

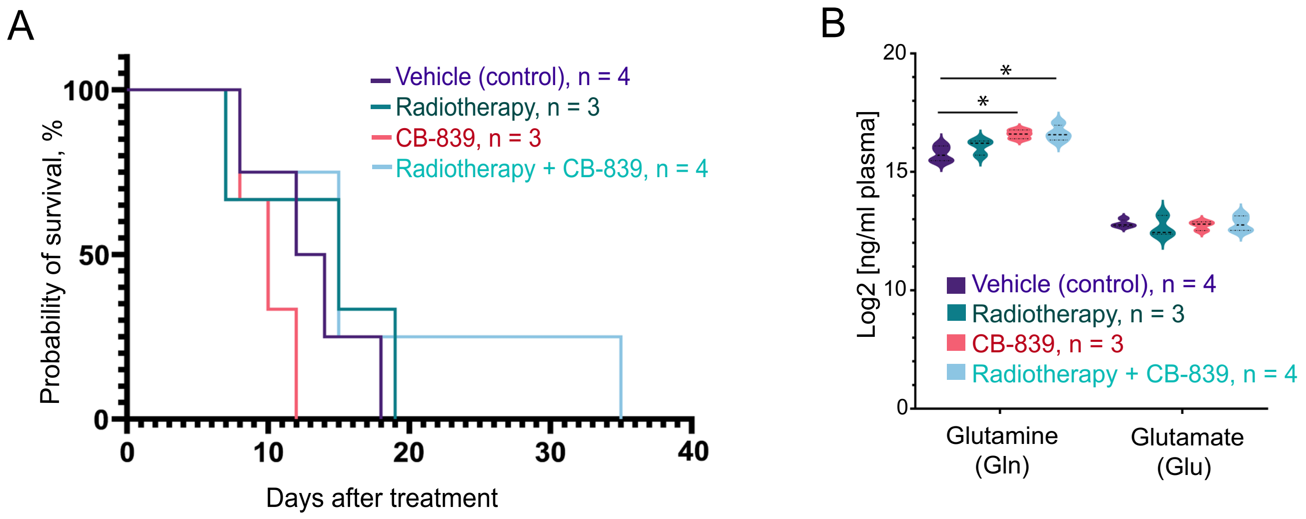

Supplementary Figure 2

A

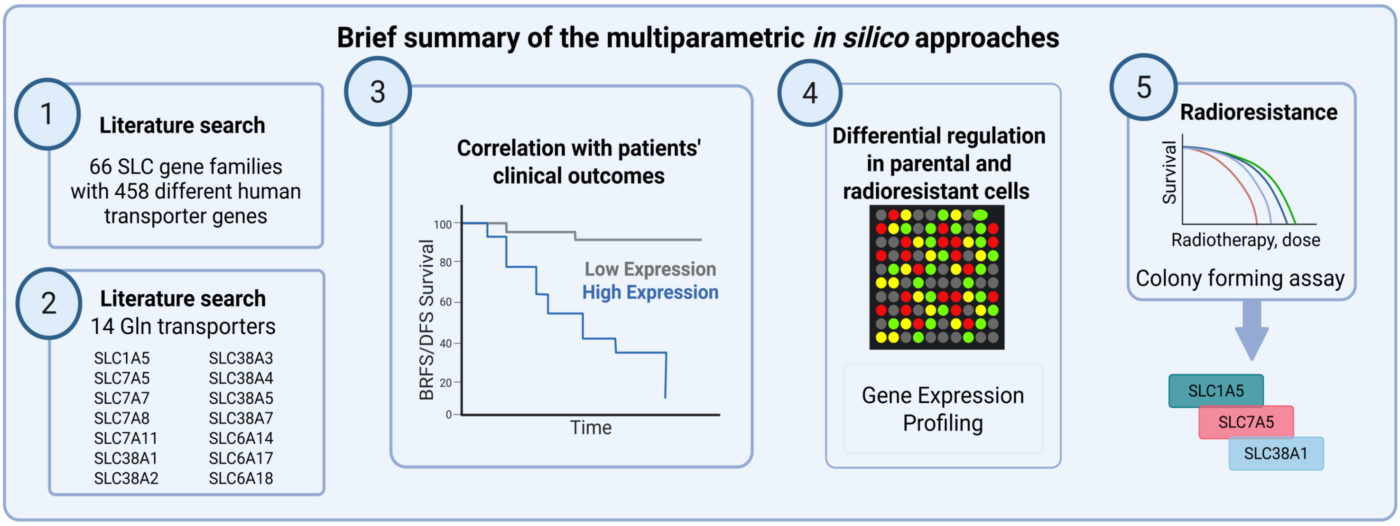


B

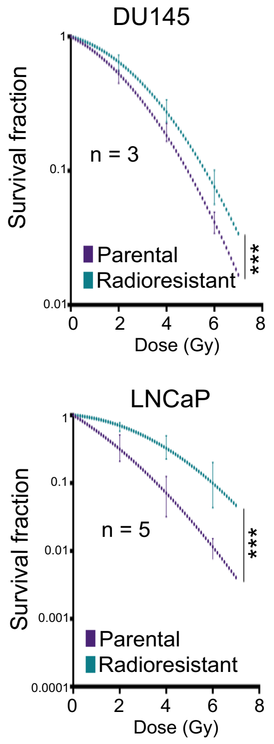


C

| Genes | TCGA PRAD | | |
|---|---|---|---|
| | OS | DFS | BRFS |
| SLC1A5 | ∅ | ⊕ | ⊕ |
| SLC38A1 | ⊕ | ⊕ | ∅ |
| SLC38A2 | ∅ | ⊕ | ∅ |
| SLC38A3 | ⊕ | ⊕ | ∅ |
| SLC38A4 | ∅ | ⊕ | ⊕ |
| SLC38A5 | ⊕ | ⊕ | ⊕ |
| SLC38A7 | ⊕ | ∅ | ∅ |
| SLC6A14 | ⊕ | ⊕ | ⊕ |
| SLC6A17 | ⊕ | ⊕ | ⊕ |
| SLC6A18 | | | |
| SLC7A11 | ⊕ | ∅ | ⊕ |
| SLC7A5 | ∅ | ∅ | ∅ |
| SLC7A7 | ∅ | ⊕ | ⊕ |
| SLC7A8 | ⊕ | ⊕ | ⊕ |

Low expression = better prognosis
High expression = better prognosis
No correlation trend
∅ No significant correlation
⊕ Statistically significant correlation

D

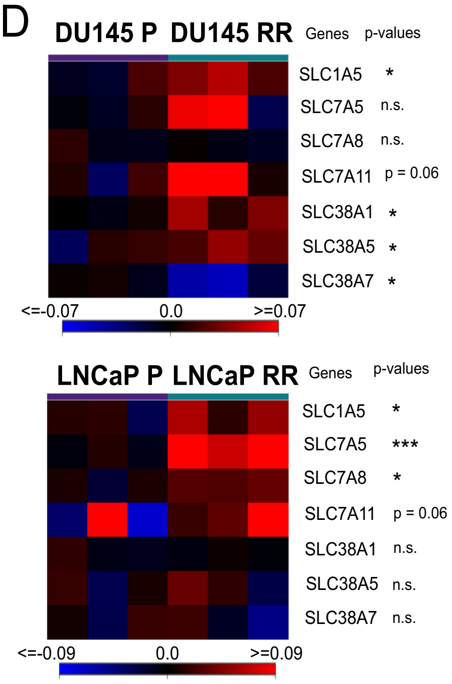

### Supplementary Figure 3

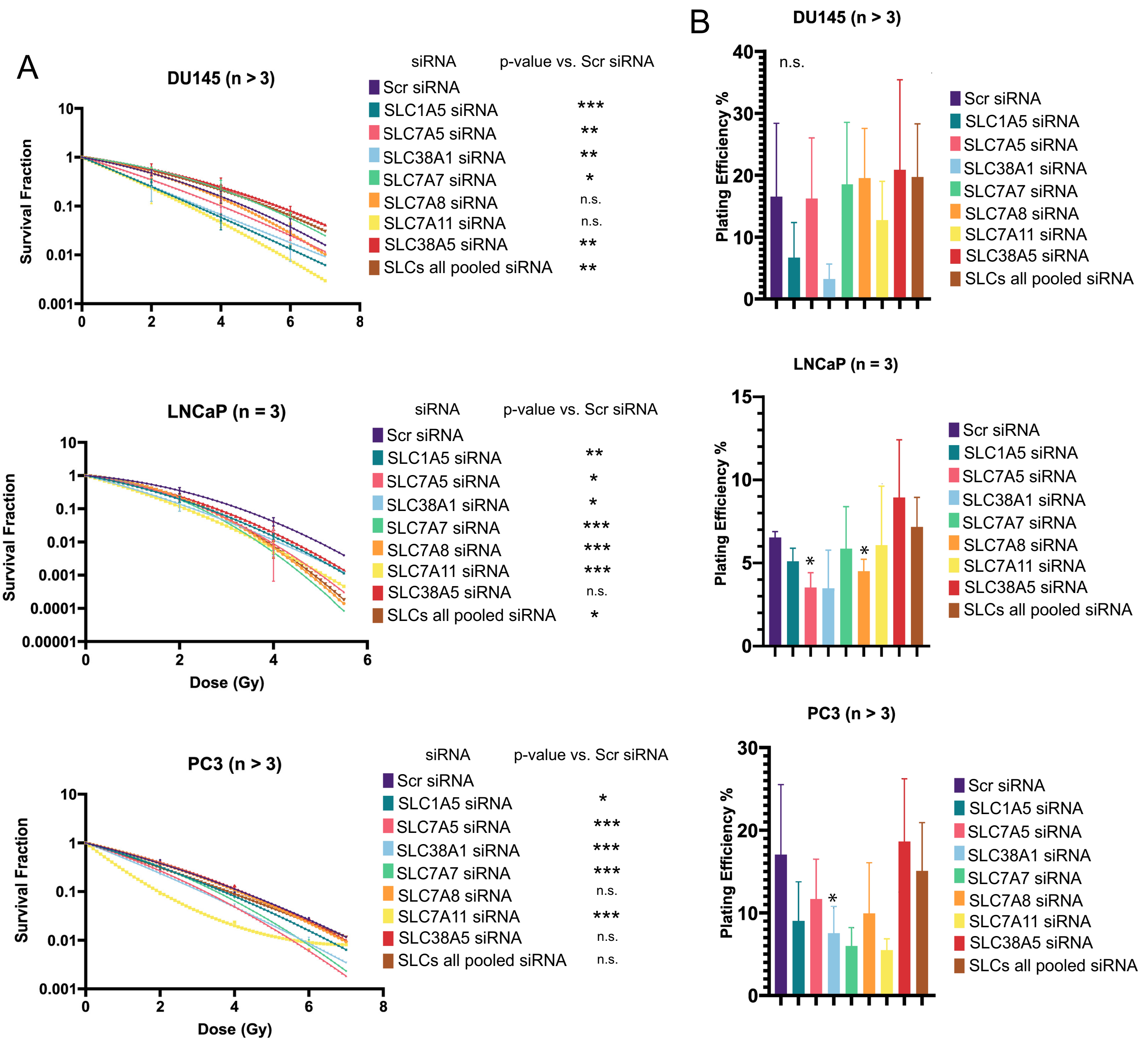

Supplementary Figure 4

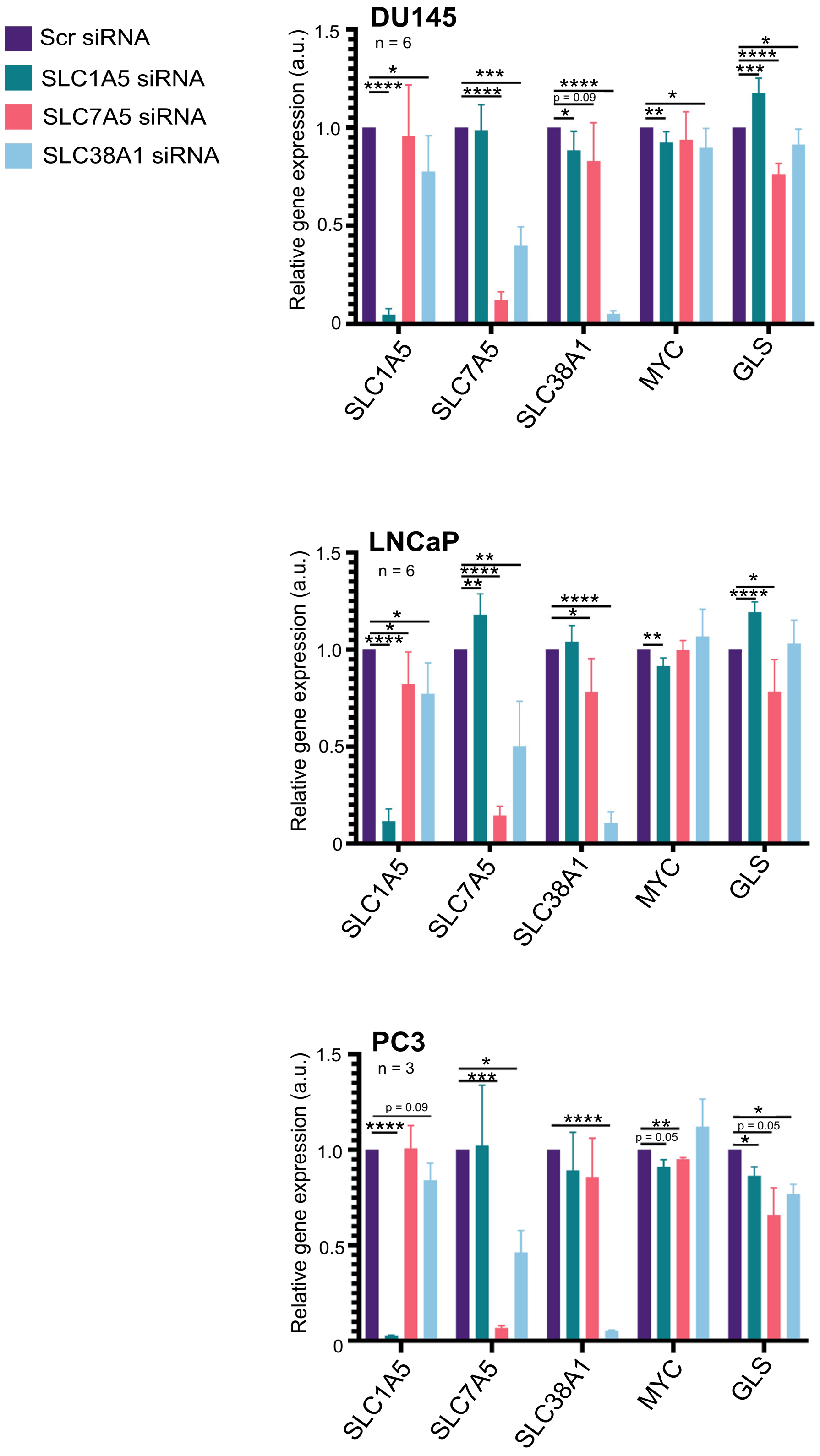

### Supplementary Figure 5

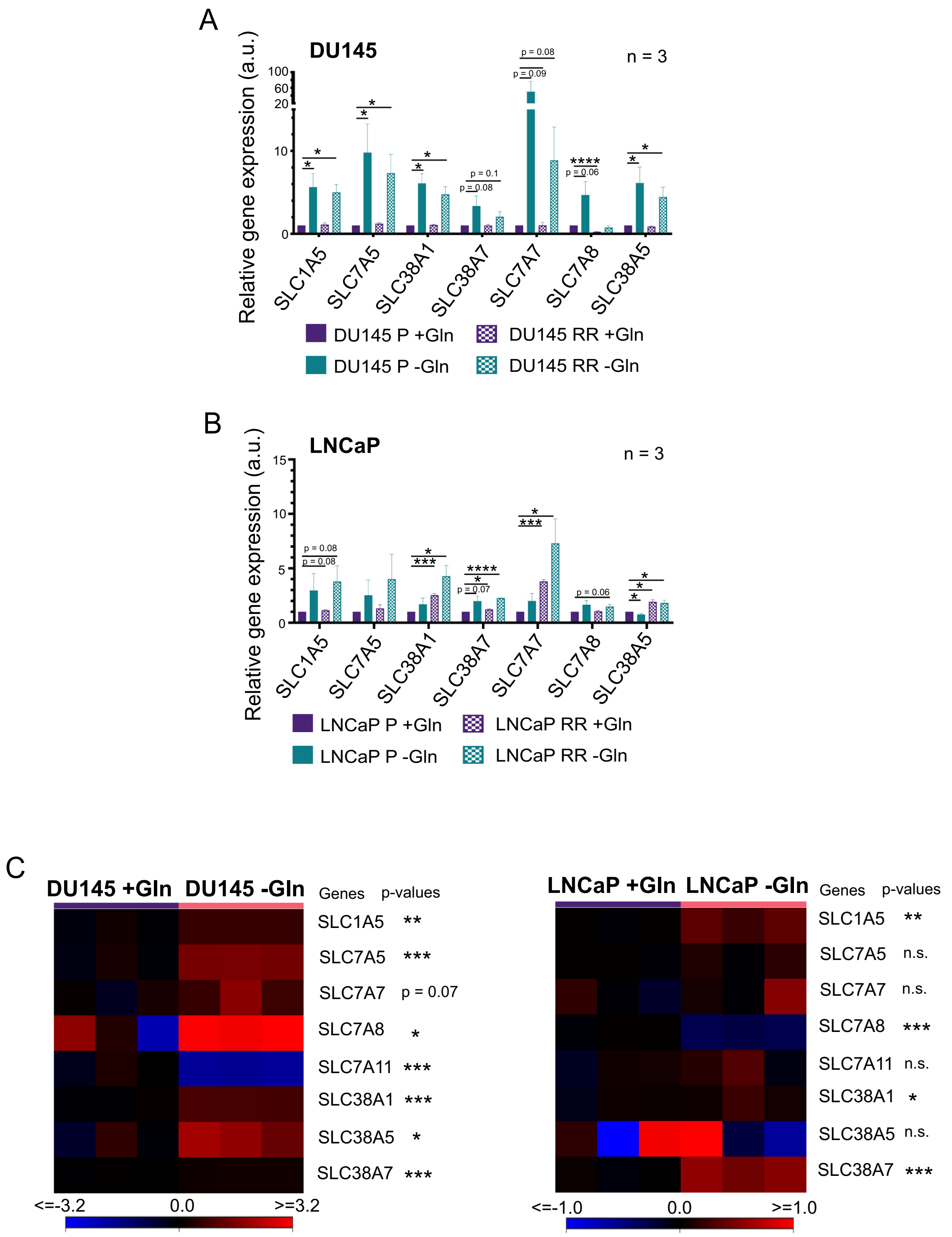

#### Supplementary Figure 6

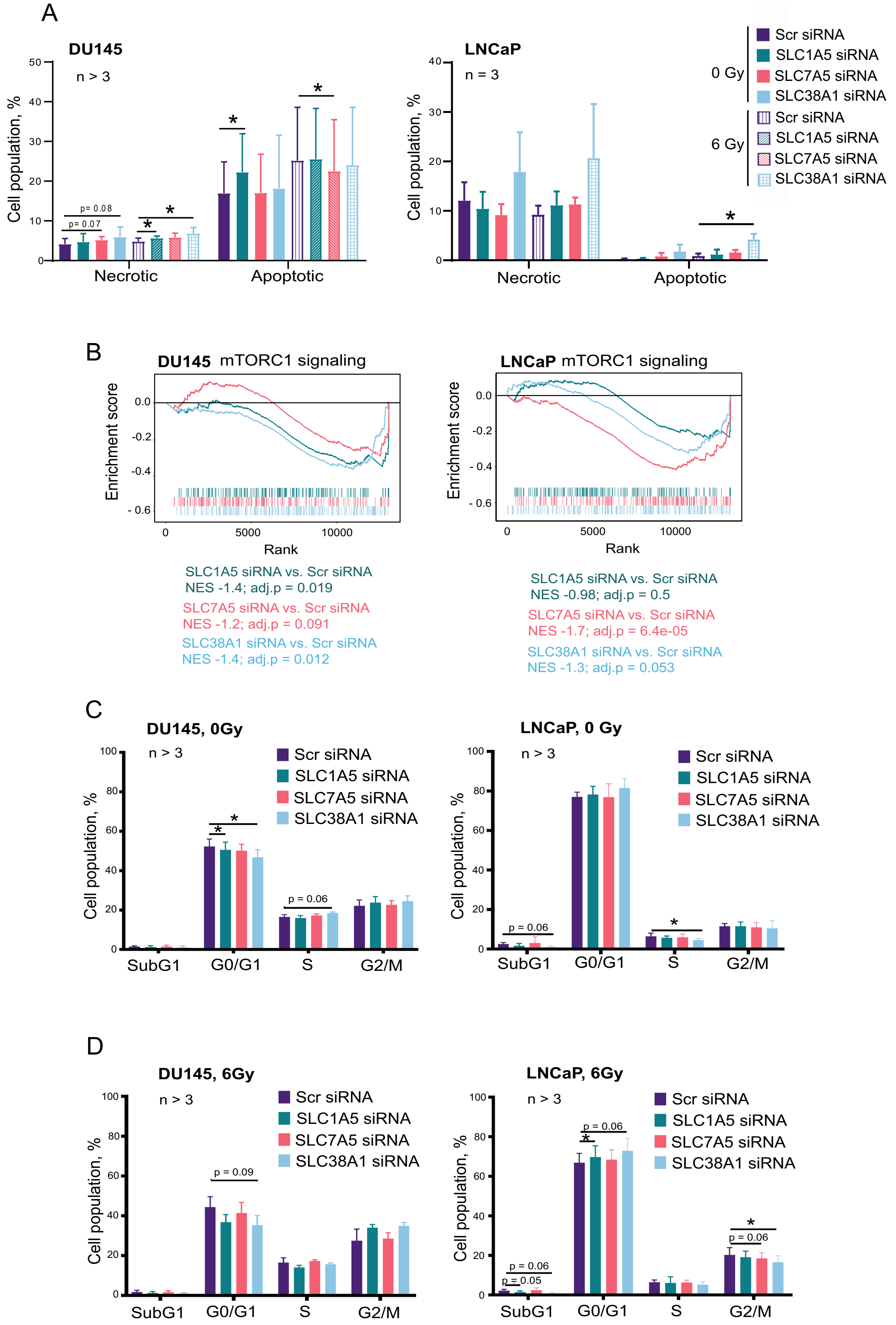

### Supplementary Figure 7

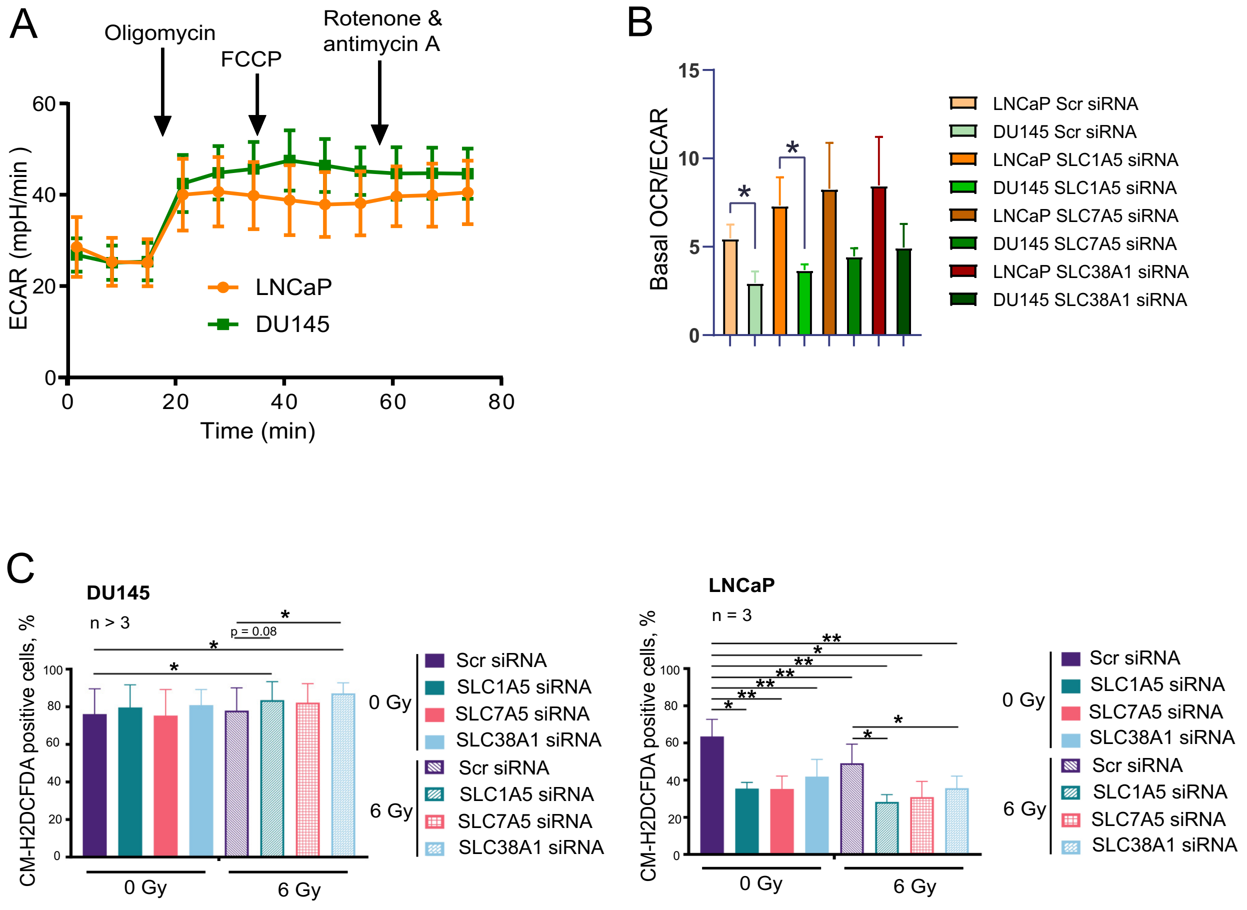

Supplementary Figure 8

A

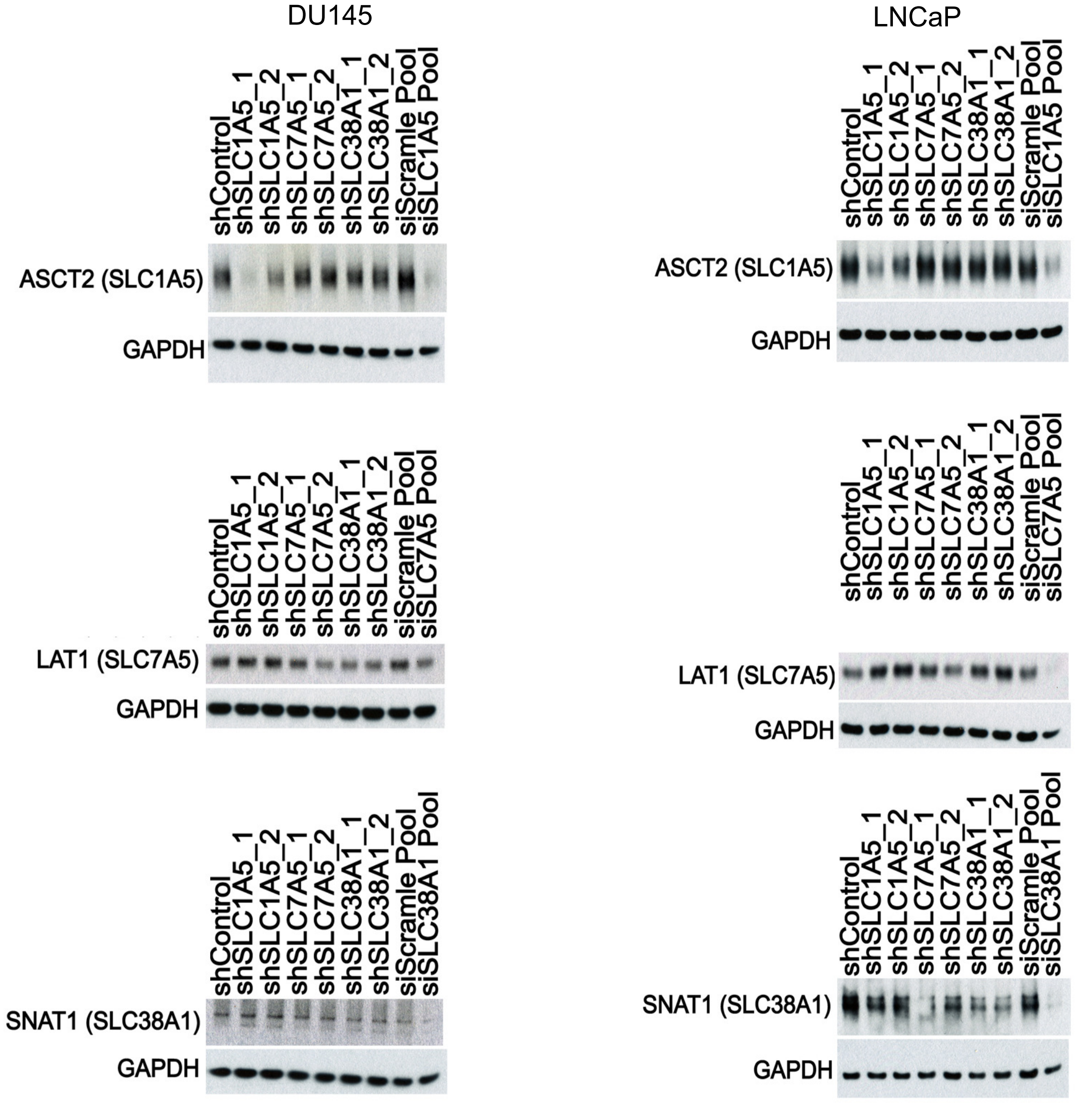


B

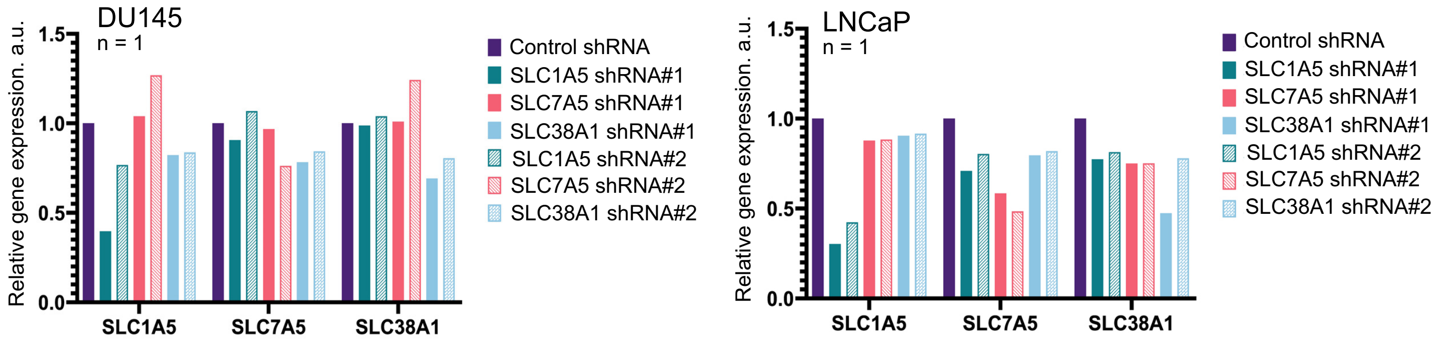

### Supplementary Figure 9

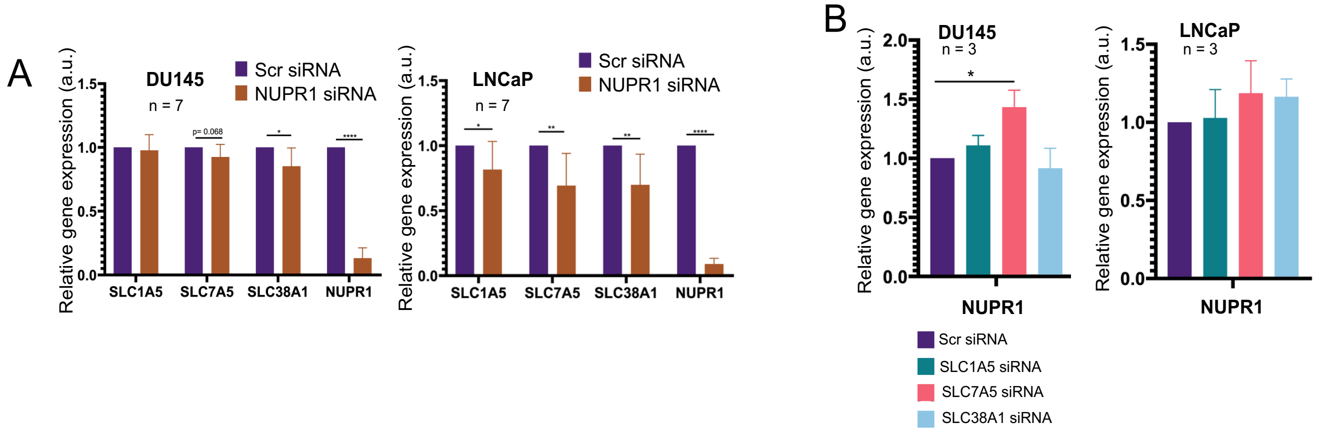


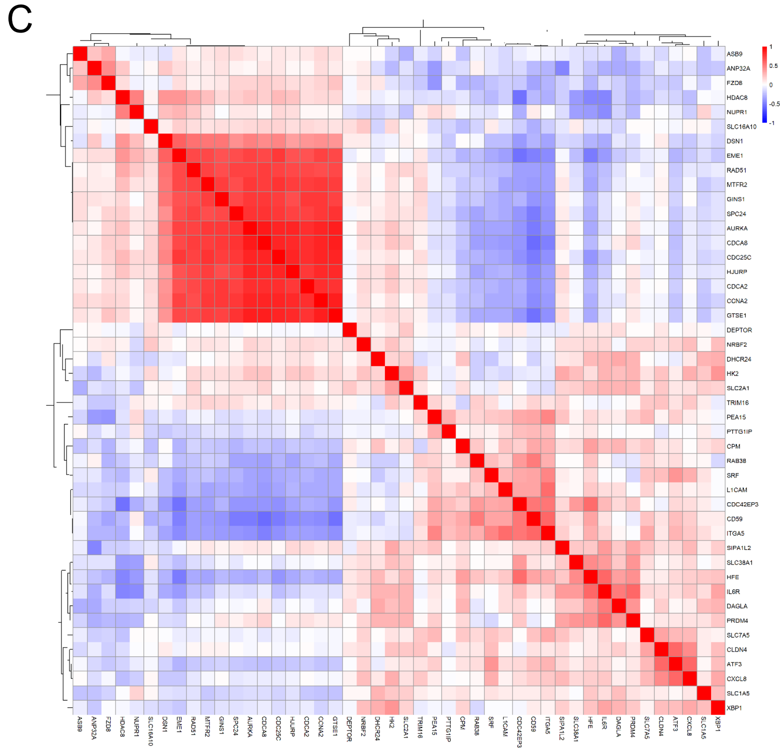


| -1 / 1 | Correlation Coefficients SLC1A5 | SLC7A5 | SLC38A1 |
|---|---|---|---|
| ANP32A | -0.22292 | -0.18701 | -0.10811 |
| ASB9 | -0.22046 | -0.07496 | -0.10741 |
| AURKA | -0.12475 | -0.03254 | -0.01775 |
| CCNA2 | -0.23436 | -0.02593 | -0.01031 |
| CDC25C | -0.17214 | -0.05454 | -0.11317 |
| CDCA2 | -0.20323 | -0.00291 | -0.0206 |
| CDCA8 | -0.13534 | -0.09787 | -0.00066 |
| DSN1 | -0.11246 | -0.00656 | -0.17931 |
| EME1 | -0.13887 | -0.06388 | -0.25615 |
| MTFR2 | -0.20437 | -0.02094 | -0.11089 |
| FZD8 | -0.17041 | -0.16774 | -0.10636 |
| GINS1 | -0.17193 | -0.02477 | -0.03999 |
| GTSE1 | -0.22486 | -0.0217 | -0.05523 |
| HDAC8 | -0.00802 | -0.23815 | -0.39732 |
| HJURP | -0.11277 | -0.06438 | -0.03383 |
| RAD51 | -0.10356 | -0.07176 | -0.13544 |
| SPC24 | -0.14675 | -0.07886 | -0.09285 |

| -1 / 1 | Correlation Coefficients SLC1A5 | SLC7A5 | SLC38A1 |
|---|---|---|---|
| ATF3 | 0.026633 | 0.263693 | 0.068877 |
| CD59 | 0.074182 | 0.2292 | 0.111082 |
| CDC42EP3 | 0.071047 | 0.002763 | 0.397881 |
| CLDN4 | 0.077274 | 0.349355 | 0.023162 |
| CPM | 0.101699 | 0.215462 | 0.146849 |
| DAGLA | 0.243696 | 0.018382 | 0.21403 |
| DEPTOR | 0.032157 | 0.01554 | 0.081661 |
| DHCR24 | 0.296877 | 0.000567 | 0.109345 |
| HFE | 0.207773 | 0.073539 | 0.447869 |
| HK2 | 0.219234 | 0.10729 | 0.247335 |
| IL6R | 0.130308 | 0.151446 | 0.381216 |
| CXCL8 | 0.088714 | 0.277026 | 0.144584 |
| ITGA5 | 0.13004 | 0.335074 | 0.061211 |
| L1CAM | 0.063228 | 0.179785 | 0.031975 |
| NRBF2 | 0.025836 | 0.160357 | 0.13406 |
| PEA15 | 0.029267 | 0.275953 | 0.056392 |
| PRDM4 | 0.112486 | 0.041324 | 0.412671 |
| PTTG1IP | 0.077618 | 0.005868 | 0.043694 |
| RAB38 | 0.036313 | 0.150404 | 0.013313 |
| SIPA1L2 | 0.071387 | 0.221945 | 0.204903 |
| SLC16A10 | 0.032144 | 0.002314 | 0.076427 |
| SLC2A1 | 0.194596 | 0.062651 | 0.197205 |
| SRF | 0.117095 | 0.2359 | 0.032339 |
| TRIM16 | 0.067784 | 0.214019 | 0.016207 |
| XBP1 | 0.256512 | 0.050651 | 0.076273 |

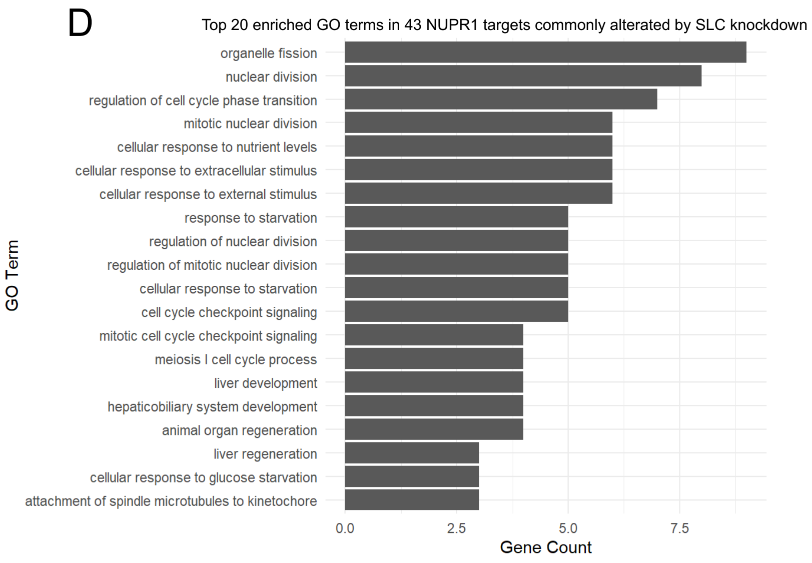

Supplementary Figure 10

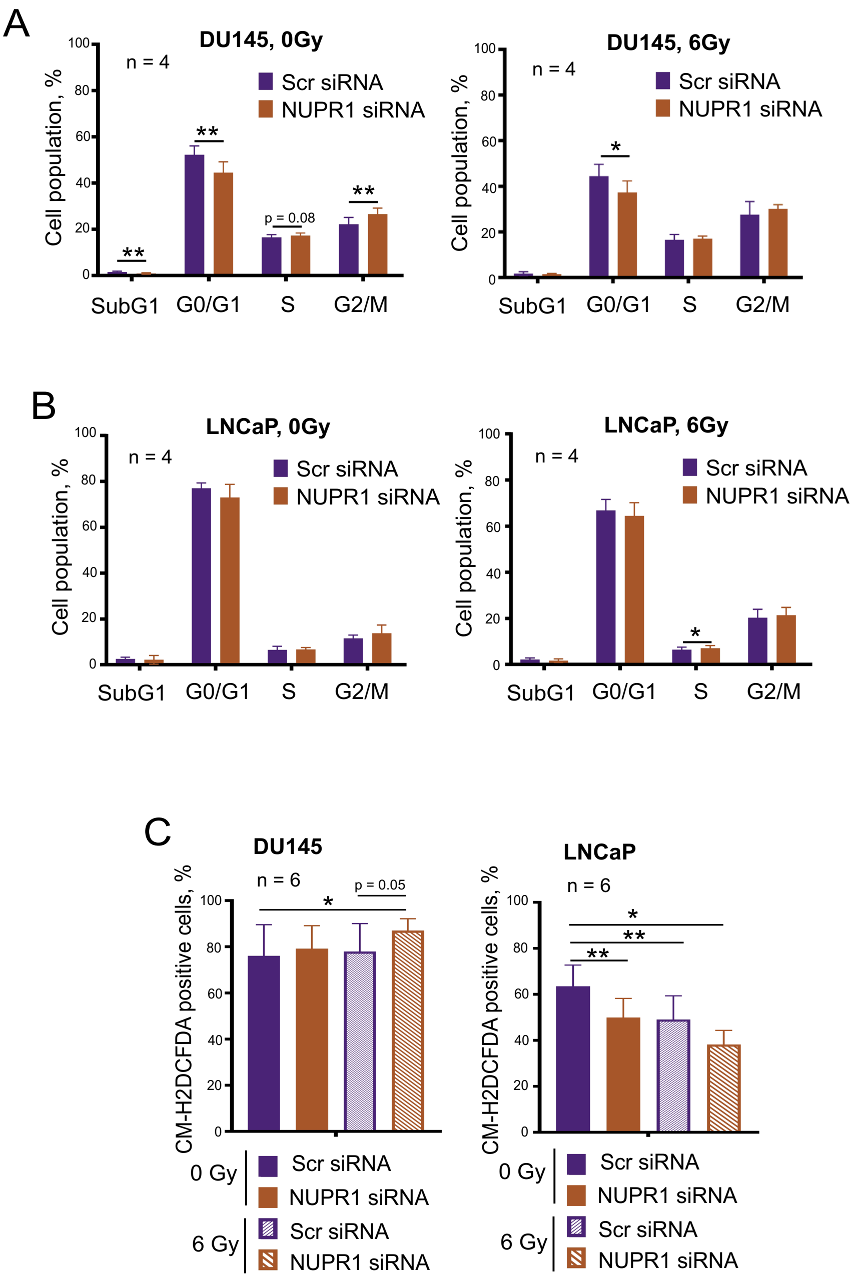

Supplementary Figure 11

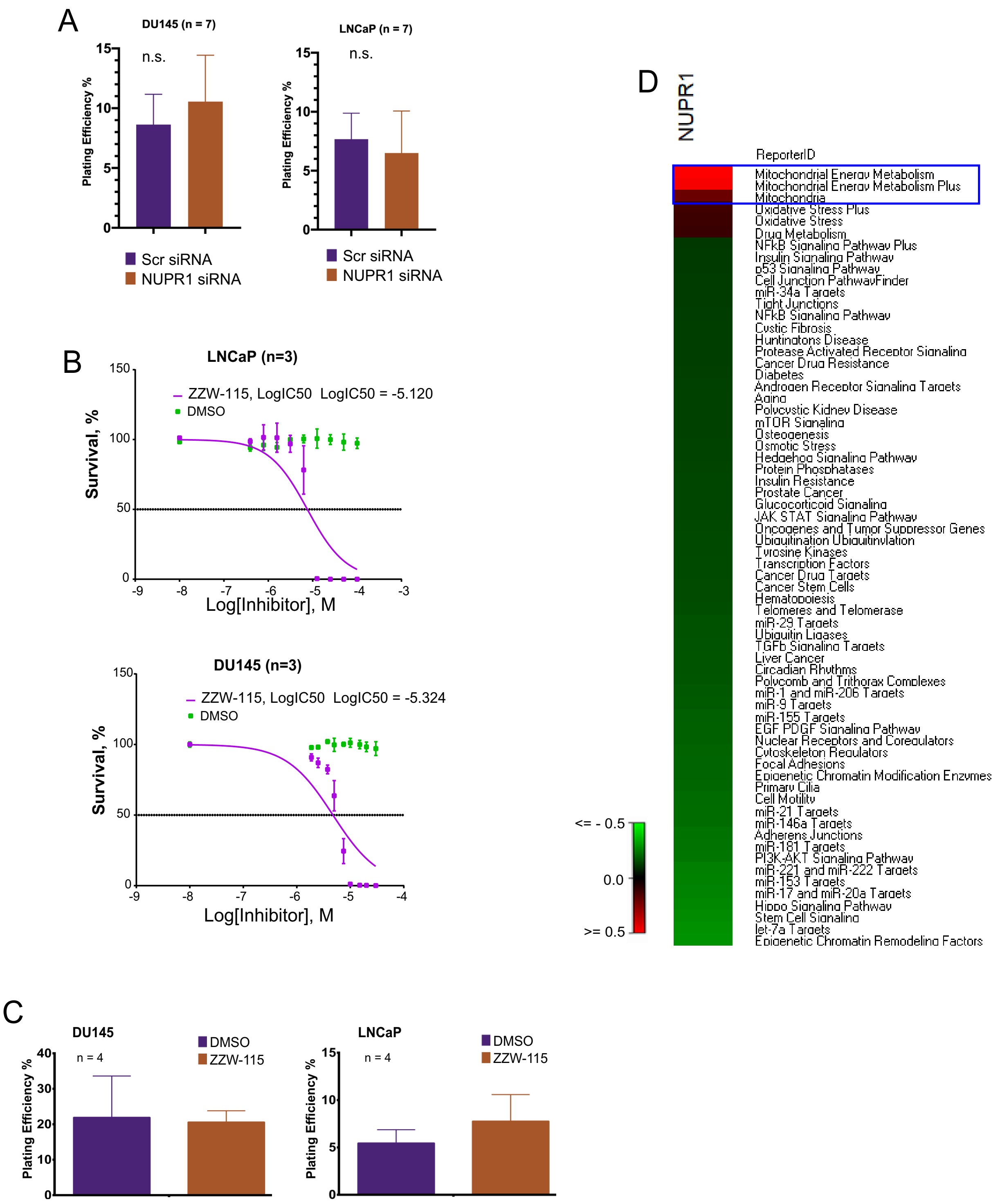

#### Supplementary references:


1. Hevler JF, Zenezeni Chiozzi R, Cabrera-Orefice A, Brandt U, Arnold S, Heck AJR: **Molecular characterization of a complex of apoptosis-inducing factor 1 with cytochrome c oxidase of the mitochondrial respiratory chain**. *Proc Natl Acad Sci U S A* 2021, **118**(39).
2. Sayyed UMH, Mahalakshmi R: **Mitochondrial protein translocation machinery: From TOM structural biogenesis to functional regulation**. *J Biol Chem* 2022, **298**(5):101870.
3. Sevrioukova IF: **Apoptosis-inducing factor: structure, function, and redox regulation**. *Antioxid Redox Signal* 2011, **14**(12):2545-2579.
4. Susin SA, Lorenzo HK, Zamzami N, Marzo I, Snow BE, Brothers GM, Mangion J, Jacotot E, Costantini P, Loeffler M *et al*: **Molecular characterization of mitochondrial apoptosis-inducing factor**. *Nature* 1999, **397**(6718):441-446.
5. Wischhof L, Scifo E, Ehninger D, Bano D: **AIFM1 beyond cell death: An overview of this OXPHOS-inducing factor in mitochondrial diseases**. *EBioMedicine* 2022, **83**:104231.
6. Rodgers JT, Lerin C, Haas W, Gygi SP, Spiegelman BM, Puigserver P: **Nutrient control of glucose homeostasis through a complex of PGC-1alpha and SIRT1**. *Nature* 2005, **434**(7029):113-118.
7. Schopf FH, Biebl MM, Buchner J: **The HSP90 chaperone machinery**. *Nat Rev Mol Cell Biol* 2017, **18**(6):345-360.
8. Myatt SS, Lam EW: **The emerging roles of forkhead box (Fox) proteins in cancer**. *Nat Rev Cancer* 2007, **7**(11):847-859.
9. Radhakrishnan SK, Gartel AL: **FOXM1: the Achilles' heel of cancer?** *Nat Rev Cancer* 2008, **8**(3):c1; author reply c2.
10. Dai C, Sampson SB: **HSF1: Guardian of Proteostasis in Cancer**. *Trends Cell Biol* 2016, **26**(1):17-28.
11. Liu L, Zhao Q, Xiong D, Li D, Du J, Huang Y, Yang Y, Chen R: **Suppressing mitochondrial inner membrane protein (IMMT) inhibits the proliferation of breast cancer cells through mitochondrial remodeling and metabolic regulation**. *Sci Rep* 2024, **14**(1):12766.
12. John GB, Shang Y, Li L, Renken C, Mannella CA, Selker JM, Rangell L, Bennett MJ, Zha J: **The mitochondrial inner membrane protein mitofilin controls cristae morphology**. *Mol Biol Cell* 2005, **16**(3):1543-1554.
13. Katiyar A, Fujimoto M, Tan K, Kurashima A, Srivastava P, Okada M, Takii R, Nakai A: **HSF1 is required for induction of mitochondrial chaperones during the mitochondrial unfolded protein response**. *FEBS Open Bio* 2020, **10**(6):1135-1148.
14. Park HJ, Carr JR, Wang Z, Nogueira V, Hay N, Tyner AL, Lau LF, Costa RH, Raychaudhuri P: **FoxM1, a critical regulator of oxidative stress during oncogenesis**. *EMBO J* 2009, **28**(19):2908-2918.
15. Black M, Arumugam P, Shukla S, Pradhan A, Ustiyan V, Milewski D, Kalinichenko VV, Kalin TV: **FOXM1 nuclear transcription factor translocates into mitochondria and inhibits oxidative phosphorylation**. *Mol Biol Cell* 2020, **31**(13):1411-1424.

**Supplementary Table 1**. The siRNA sequences used in the study.

| siRNAs | siRNA ID | Target sequence (5'-3') |
|---|---|---|
| Scr Pool | ScrambledMCR | GCAGCUAUAUGAAUGUUGU |
| | NonSpecific68%GC | UGCGCUAGGCCUCGGUUGC |
| | Scr EGFP | GGCUACGUCCAGGAGCGCA |
| SLC7A5 Pool | SLC7A5 #1 | GAGGAUGGAAUUACUUGAA |
| | SLC7A5 #2 | GCCUCAAGGAUACAGGGAG |
| | SLC7A5 #3 | AACAGAGACAAGAAAGGCA |
| SLC1A5 Pool | SLC1A5-1260 | GGAAUAUCACCGGAACCAG |
| | SLC1A5-896 | CCUCAAUGUAGAAGGUGAC |
| | SLC1A5-948 | GACCGUACGGAGUCGAGAA |
| SLC38A1 Pool | SLC38A1-130944 | CCGAAGUAGAAAAUGGUCA |
| | SLC38A1-HSS129873 | CAGUGCCCGAGGAUGAUAA |
| | SLC38A1-s37590 | GGAGUUACAUCUGCUAACA |
| SLC7A8 Pool | SLC7A8-1909 | GCGACAUGUACACACUCAU |
| | SLC7A8-2314 | CCACGAAGGACAAGGACGU |
| | SLC7A8-2212 | AGAUGUGUGUGGUCGUGUA |
| SLC38A5 Pool | SLC38A5-1 | CGGGAUAUCUUUGGAGUUA |
| | SLC38A5-2 | CUUCGGAUGCUGUGGGCUA |
| | SLC38A5-3 | GCUACAGGCAAGAACGUGA |
| | SLC38A5-4 | GCCGGUCCAGUUCAUGGAU |
| SLC7A7 Pool | SLC7A7-161 | GGUUGACAGCACUGAGUAU |
| | SLC7A7-997 | CCUAUUAUACUGUGCUAGA |
| | SLC7A7-416 | GGCCCUUUGUUAUGCGGAA |
| | SLC7A7-1635 | GAACAUAAGCGACCGCUUU |
| SLC7A11 Pool | SLC7A11-5167 | ACAUAUCAAUAGAGGGUUA |
| | SLC7A11-46 | GGAACGAGGAGGUGGAGAA |
| | SLC7A11-3060 | AAUGAGAAUCUGUGGAUAA |
| | SLC7A11-1511 | CGAUACAAAUGCCCAGAUA |
| NUPR1 Pool | NUPR1-253 | AGAGGAAACUGGUGACCAA |
| | NUPR1-473 | GAAAGCGCCUCCAACCCUA |
| | NUPR1-355 | CAGCAAUAGAGACGGGACU |
| | NUPR1-187 | GCCGGAAAGGUCGCACCAA |
| MYC Pool | MYC-23 | ACGGAACUCUUGUGCGUAA |
| | MYC-24 | GAACACACAACGUCUUGGA |
| | MYC-25 | AACGUUAGCUUCACCAACA |
| | MYC-26 | CGAUGUUGUUUCUGUGGAA |
| GLS Pool | GLS-671 | GGGUCUGUUACCUAGCUUG |
| | GLS-720 | GGACAAGAGAAAAUACCUG |
| | GLS-1626 | GCAGUUCGAAAUACAUUGA |

**Supplementary Table 2.** The shRNA expression vectors used in the study.

| shRNAs | shRNA ID | Clone ID | Vector Backbone | Target Sequences |
|---|---|---|---|---|
| shSLC1A5 | shSLC1A5-1 | TRCN0000043118 | pLKO.1 | CTGGATTATGAGGAATGGATA |
| | shSLC1A5-2 | TRCN0000043120 | pLKO.1 | GCCTGAGTTGATACAAGTGAA |
| shSLC7A5 | shSLC7A5-1 | TRCN0000043008 | pLKO.1 | GCATTATACAGCGGCCTCTTT |
| | shSLC7A5-2 | TRCN0000043009 | pLKO.1 | CTAGATCCCAACTTCTCATTT |
| shSLC38A1 | shSLC38A1-1 | TRCN0000043900 | pLKO.1 | CCTCCTATTGATCTGTTCAAA |
| | shSLC38A1-2 | TRCN0000043901 | pLKO.1 | CCTGCATTGTTCCAGAGCTAA |
| shControl | pLKO.1 | | pLKO.1 | CCTAAGGTTAAGTCGCCCTCG |

**Supplementary Table 3.** The primers used in the study.

| | | |
|---|---|---|
| SLC1A5 | SLC1A5-Fw2 | CACCATGGTTCTGGTCTCCTG |
| | SLC1A5-Rv2 | GGTGAAGAGGAAGTAGATGAGG |
| SLC7A5 | SLC7A5-Fw | CCTTCATCGCAGTACATCGTGG |
| | SLC7A5-Rv | GCCTTCACGCTGTAGCAGTTCAC |
| SLC38A1 | SLC38A1-Fw | GCCATTATGGGCAGTGGGAT |
| | SLC38A1-Rv | ACACCATGCAGCCTGTTTCT |
| MYC | c-MYC-Fw | CTCCGTCCTCGGATTCTCTGC |
| | c-MYC-Rv | CTCCAGCAGAAGGTGATCCAG |
| GLS | GLS1-Fw | GAGGCATTCTACTGGAGATACC |
| | GLS1-Rev | GCTCCAGCATTTACCATAGG |
| SLC7A7 | SLC7A7-Fw | CGAGCTGCTTCGCCCCTTATG |
| | SLC7A7-Rv | TGACCGCGATCAGTGCCAATAC |
| SLC7A8 | SLC7A8-Fw | CCACATTTGGAGGAGTTAATGG |
| | SLC7A8-Rv | GTACATGTCGCTGGTGACC |
| SLC7A11 | SLC7A11-Fw | GCGTGGGCATGTCTCTGAC |
| | SLC7A11-Rv | GCTGGTAATGGACCAAAGACTTC |
| SLC38A5 | SLC38A5-Fw | CCTGGTTATCGGCACCTTCC |
| | SLC38A5-Rv | CCAAGTGTTTCATGAGGGCGA |
| SLC38A7 | SLC38A7-Fw | CGACCAGCAGGACAAGATTAT |
| | SLC38A7-Rv | GATGAAGAGGAAGGCAGTGAG |
| NUPR1 | NUPR1-Fw | GCACCAAGAGAGAAGCTGC |
| | NUPR1-Rv | GGTCTGGCCTCATCTCCAG |
| ATF4 | ATF4-Fw | CCAACAACAGCAAGGAGGATGC |
| | ATF4-Rv | GACTAGGGGGGCAAAGAGATCAC |
| EZH2 | EZH2_for | GAgTGTTCGGTGACCAGTGA |
| | EZH2_rev | CCCGTGTACTTTCCCATCAT |
| BMI1 | BMI1 Fw | GAGGTTGCAGATGAAGATAAGAG |
| | BMI1 Rv | CTGGAAAGTATTAGGTATGTCC |
| ACTB | ACTB-Fw | ATGGAGTCCTGTGGCATCCA |
| | ACTB-Rev | AGTACTTGCGCTCAGGAGGA |
| RPLP0 | RPLP0-Fw | CTCAACATCTCCCCCTTCTCCTT |
| | RPLP0-Rev | TGATGCAACAGTTGGGTAGCC |

**Supplementary Table 4.** Antibodies used for Western blot analysis.

| Antibodies | Catalog Number | Name | Dilution | Company |
|---|---|---|---|---|
| SLC1A5 | #8057 | ASCT2 (D7C12) Rabbit mAb | 1:1000 | Cell Signaling Technology |
| SLC7A5 | sc-374232 | LAT1 (D-10) | 1:1000 | Santa Cruz Biotechnology |
| SLC38A1 | #36057 | SNAT1/SLC38A1 (D9L2P) Rabbit mAb | 1:1000 | Cell Signaling Technology |
| GAPDH | #sc-25778 | GAPDH (FL-335) | 1:1000 | Santa Cruz Biotechnology |
| γH2A.X (Ser139) | # 05-636 | Anti-phospho-Histone H2A.X (Ser139) | 1:1000 | Sigma-Aldrich |
| Anti-Rabbit IgG (Alexa Fluor 488) | #A32731 | Anti-Rabbit IgG (Alexa Fluor 488), Donkey | 1:350 | Thermo Fisher Scientific |

**Supplementary Table 5.** A comparative phosphoproteomics analysis of PCa cell lines DU145 and LNCaP with or without knockdown of Gln AATs.

| Uniprot ID | Entrez Gene | Gene Symbol | **log2 DU145 shSCR / DU145 shSLC1A5_1** | **log2 DU145 shSCR / DU145 shSLC7A5_2** | **log2 DU145 shSCR / DU145 shSLC38A1_1** | **pVal** |
|---|---|---|---|---|---|---|
| Q15388 | 9804 | TOMM20 | 1.115477 | 0.891851 | 0.606179 | 0.004101 |
| A0A6Q8PFE1 | 9131 | AIFM1 | 0.315573 | 0.312384 | 0.396087 | 0.000238 |
| Q56NI9 | 157570 | ESCO2 | 0.487969 | 0.485427 | 0.393504 | 0.000126 |
| P07900 | 3320 | HSP90AA1 | -0.57167 | -0.55439 | -0.71209 | 0.000253 |
| Q96EB6 | 23411 | SIRT1 | -0.31171 | -0.36355 | -0.49228 | 0.001921 |

| Uniprot ID | Entrez Gene | Gene Symbol | **log2 LNCaP shSCR / LNCaP shSLC1A5_1** | **log2 LNCaP shSCR / LNCaP shSLC7A5_2** | **log2 LNCaP shSCR / LNCaP shSLC38A1_1** | **pVal** |
|---|---|---|---|---|---|---|
| A0A2P9DTZ8 | 2305 | FOXM1 | 0.329965 | 0.440997 | 0.593 | 0.004101255 |
| Q00613 | 3297 | HSF1 | 0.310598 | 0.393149 | 0.297655 | 0.000238236 |
| Q08999 | 5934 | RBL2 | 0.393664 | 0.590192 | 0.672692 | 0.644468847 |
| Q05513 | 5590 | PRKCZ | 0.411925 | 0.311944 | 0.405679 | 0.003250252 |
| A0A075B6F9 | 51070 | NOSIP | 0.407897 | 0.686894 | 0.915098 | 0.000253379 |
| Q16891 | 10989 | IMMT | 0.767339 | 0.553936 | 0.67423 | 0.001921349 |
| Q4LE36 | 47 | ACLY | 0.351763 | 0.453547 | 0.822654 | 0.004101255 |
| Q13057 | 80347 | COASY | 0.364851 | 0.332103 | 0.375552 | 0.000238236 |

**Supplementary Table 6.** The upstream regulators identified in PCa cell lines DU145 and LNCaP using Ingenuity Pathway Analysis (IPA) in response to specified treatments.

| DU145 siSLC1A5 | | DU145 siSLC7A5 | | DU145 siSLC38A1 | | LNCaP siSLC1A5 | | LNCaP siSLC7A5 | | LNCaP siSLC38A1 | |
|---|---|---|---|---|---|---|---|---|---|---|---|
| Upstream Regulator | p-value | Upstream Regulator | p-value | Upstream Regulator | p-value | Upstream Regulator | p-value | Upstream Regulator | p-value | Upstream Regulator | p-value |
| SYVN1 | 4.12E-08 | ERBB2 | 1.83E-06 | NUPR1 | 1.83E-12 | SKP2 | 5.37E-05 | CCND1 | 5.58E-07 | E2F4 | 6.66E-26 |
| ERBB2 | 2.68E-06 | COPS5 | 2.45E-06 | SUZ12 | 1.20E-11 | FLOT1 | 6.67E-05 | let-7 | 1.18E-06 | NUPR1 | 4.15E-24 |
| TCF7L2 | 7.92E-06 | GLI1 | 2.77E-06 | ERBB2 | 7.77E-10 | TRPC1 | 6.80E-05 | NUPR1 | 1.67E-06 | ERBB2 | 1.47E-22 |
| NUPR1 | 1.62E-05 | NUPR1 | 9.72E-06 | CDK4 | 2.14E-08 | PRDM4 | 8.56E-05 | SUZ12 | 9.40E-06 | CCND1 | 5.06E-18 |
| SUZ12 | 8.18E-05 | SYVN1 | 1.49E-05 | NR3C1 | 1.09E-07 | NUPR1 | 1.38E-04 | CDK4 | 1.18E-05 | RABL6 | 4.14E-17 |

| DU145 P +/- Gln | | DU145 RR +/- Gln | | LNCaP P +/- Gln | | LNCaP RR +/- Gln | |
|---|---|---|---|---|---|---|---|
| Upstream Regulator | p-value | Upstream Regulator | p-value | Upstream Regulator | p-value | Upstream Regulator | p-value |
| NUPR1 | 7.31E-32 | NUPR1 | 5.61E-24 | E2F4 | 2.98E-31 | NUPR1 | 2.51E-09 |
| TP53 | 5.52E-20 | TP53 | 1.17E-13 | TBX2 | 1.47E-26 | GNE | 1.43E-08 |
| TGFB1 | 6.57E-16 | ERBB2 | 1.82E-12 | NUPR1 | 1.93E-26 | CCND1 | 2.05E-08 |
| ERBB2 | 1.64E-15 | ESR1 | 1.17E-11 | RABL6 | 3.08E-26 | tosedostat | 2.25E-07 |
| ESR1 | 7.65E-14 | RABL6 | 7.09E-11 | CCND1 | 2.65E-25 | FOXM1 | 1.51E-06 |
